# Occipital white matter tracts in human and macaque

**DOI:** 10.1101/069443

**Authors:** Hiromasa Takemura, Franco Pestilli, Kevin S. Weiner, Georgios A. Keliris, Sofia M. Landi, Julia Sliwa, Frank Q. Ye, Michael A. Barnett, David A. Leopold, Winrich A. Freiwald, Nikos K. Logothetis, Brian A. Wandell

## Abstract

We compare the major white matter tracts in human and macaque occipital lobe using diffusion MRI. The comparison suggests similarities but also significant differences in spatial arrangement and relative sizes of the tracts. There are several apparently homologous tracts in the two species, including the vertical occipital fasciculus (VOF), optic radiation, forceps major, and inferior longitudinal fasciculus (ILF). There is one large human tract, the inferior fronto-occipital fasciculus, with no corresponding fasciculus in macaque. The macaque VOF is compact and its fibers intertwine with the dorsal segment of the ILF, but the human VOF is much more elongated in the anterior-posterior direction and may be lateral to the ILF. These similarities and differences will be useful in establishing which circuitry in the macaque can serve as an accurate model for human visual cortex.

**Contact information:** Hiromasa Takemura, Center for Information and Neural Networks (CiNet), National Institute of Information and Communications Technology, and Osaka University, Japan

**Author contribution:** Designed the study: HT FP BAW. Performed the experiments. HT FP GAK SML JS FQY DAL WAF NKL. Analyzed the data. HT FP KSW MAB BAW. Contributed analysis tools. FP KSW MAB. Wrote the paper. HT FP KSW BAW.

## Introduction

The macaque monkey has been an important model for understanding human vision. The two species perform at similar levels on many basic sensory tasks, such as color, motion and spatial discriminations (De Valois and Jacobs 1968; De Valois et al. 1974; Newsome et al. 1989; Miura et al. 2006; Horwitz 2015), or categorization of objects (Rajalingham et al. 2015). Some performance differences between human and macaque vision have also been described (Gellman et al. 1990; Zarco et al. 2009; Lindbloom-Brown et al. 2014; Horwitz 2015). A substantial literature compares human and macaque functional cortical responses to visual stimuli (Brewer et al. 2002; Tsao et al. 2003, 2008; Sasaki et al. 2006; Kriegeskorte et al. 2008; Wade et al. 2008; Pinsk et al. 2009; Mantini et al. 2012; Okazawa et al. 2012; Polosecki et al. 2013; Goda et al. 2014; Kolster et al. 2014; Russ and Leopold 2015; Lafer-Sousa et al. 2016). There are similarities and differences in the functional responses, as well (Tootell et al. 2003; Wandell and Winawer 2011; Vanduffel et al. 2014).

The anatomical connections to visual cortex are an important source for understanding the similarities and differences. It is well established that the physiological properties of individual neurons are similar throughout cortex, with large variations in neuronal responses arising substantially from differences in their connections. For instance, Wernicke’s classic theory of Connectionism emphasizes the important role of the white matter axon bundles (fascicles, tracts) in defining the long-range neuronal connections (Catani and Ffytche 2005). These tracts establish critical features of the brain’s information processing (Bullock et al. 2005; Fields 2008a, 2008b, 2015; Wandell and Yeatman 2013). Comparing the anatomical connections in the two species may help us specify which macaque behaviors and responses are a good model for human and which are not.

Diffusion MRI (dMRI) and tractography algorithms provide an excellent opportunity to better understand the organization of major white matter tracts (Catani et al. 2002; Mori and Zhang 2006; Schmahmann et al. 2007; Catani and Thiebaut de Schotten 2012; Craddock et al. 2013; Wandell and Yeatman 2013; Wandell 2016). These methods assess the large-scale organization of the major white matter tracts, including their relative positions of the tracts, their relative sizes, and the positions of the estimated tract endpoints with respect to the visual field maps (Greenberg et al. 2012; Takemura et al. 2016b). The goal of this study is to shed light on how the pattern of long-range white matter tracts contribute to the functional and behavioral similarities and differences between human and macaque visual system. This work is part of a growing literature using dMRI and tractography to compare the white matter tracts between the two species (Schmahmann et al. 2007; Rilling et al. 2008; Thiebaut de Schotten et al. 2011; Jbabdi et al. 2013; Li et al. 2013; Mars et al. 2015; van den Heuvel et al. 2016).

We report three principal findings. First, several white matter tracts are homologous in the human and macaque occipital lobe. These are the Optic Radiation (OR), Forceps Major, and Inferior Longitudinal Fasciculus (ILF). In contrast, we do not find support for the existence of a macaque tract homologous to the human Inferior Fronto-Occipital Fasciculus (IFOF), a relatively large tract in human.

Second, we describe the Vertical Occipital Fasciculus (VOF), a major white matter tract communicating between the dorsal and ventral streams (Takemura et al. 2016b). We focus on the VOF because other major tracts have been extensively described (Schmahmann et al. 2007), but until recently the VOF has been relatively neglected in both the human and macaque literature (Yeatman et al. 2013, 2014; Duan et al. 2015; Takemura et al. 2016b; Weiner et al., 2016a). The diffusion measurements find a tract located in a position consistent with classical reports by Wernicke in an unnamed monkey species (Wernicke 1881). We estimated that the dorsal and ventral endpoints of the VOF are near similar visual field maps in human and macaque. The VOF properties may be particularly important for understanding how information is communicated between the dorsal and ventral streams within the occipital lobe.

Third, we describe a difference in the relative position of several, large tracts in the occipital lobe, VOF, ILF, and OR. In both human and macaque, the VOF is clearly located lateral to the OR. However, the macaque VOF intermingles with the dorsal segment of the ILF, while the human VOF is more separated from human ILF in terms of tract position and endpoints.

## Materials and Methods

The analyses are based on a diverse set of macaque and human measurements, pooled across several laboratories and public datasets. The datasets have different spatial and angular resolution and image quality.

### MR data acquisition

#### Macaque diffusion data 1 (subject M1, *ex vivo*)

This dataset was acquired from a post-mortem *Macaca mulatta* (rhesus macaque monkey) brain using 30-cm magnet bore Bruker 7T scanner at the National Institute of Health. The dMRI data were sampled in 121 directions at a spatial resolution of 250 µm isotropic. The b-value was set to 4800 s/mm^2^ and TE was 34 ms. Seven non-diffusion weighted image (b=0) were obtained. These data were analyzed in two previous publications (Thomas et al. 2014; Reveley et al. 2015).

#### Macaque diffusion data 2 (subject M2, *in vivo*)

This dataset was acquired from a living *Macaca mulatta* (rhesus macaque monkey) brain using a Bruker BIOSPEC 4.7 T vertical bore scanner at Max Planck Institute, Tübingen, Germany. The dMRI data were sampled in 61 different directions at a spatial resolution of 750 µm isotropic.

The gradient b-value was 1200 s/mm^2^ and TE was 62.65 msec. Seven non-diffusion weighted images (b=0) were acquired at the beginning of the scan. The total imaging time was 6 h 48 min. T1-weighted images at 375 um isotropic resolution were also acquired using a 3D-MDEFT sequence (Lee et al. 1995).

#### Macaque diffusion data 3 (subject M3, *ex vivo*)

This dataset was acquired in a formalin-fixed post-mortem *Macaca mulatta* (rhesus macaque monkey) brain using a Bruker BIOSPEC 4.7 T vertical bore scanner at Max Planck Institute, Tübingen, Germany. The brain was perfused, removed from the skull, and kept in 4% paraformaldehyde for 5 years prior to scanning. The dMRI data were sampled in 61 directions at a spatial resolution of 800 µm isotropic. The b-value was 4000 s/mm^2^ and TE was 78 msec.

Seven non-diffusion weighted images (b=0) were acquired at the beginning of the scan. These measurements were repeated 4 times and then averaged. The total imaging time was 64 hours. These data were analyzed in a previous publication (Iturria-Medina et al. 2011).

#### Macaque diffusion data 4 (subject M4, *in vivo*)

This dataset was acquired from a living *Macaca mulatta* (rhesus macaque monkey) brain using a 3T SIEMENS Trio MRI scanner using an AC-88 gradient insert (Siemens) at the Citigroup Biomedical Imaging Center of the Weill Cornell Medical College. The dMRI data were sampled in 64 directions at a spatial resolution of 1 mm isotropic. The b-value was set to 2000 s/mm^2^ and TE was 76.8 msec. The measurements were repeated 3 times. A total of 42 non-diffusion weighted images (b=0) were acquired at the beginning of the session and between scans.

#### Human diffusion data: HCP90 dataset (subjects H1-H3)

We used human diffusion-weighted MRI dataset acquired from three subjects by the Human Connectome Consortium (HCP90 dataset; Van Essen et al. 2013). The HCP90 dataset was acquired at multiple b-values (1000, 2000 and 3000 s/mm^2^) and spatial resolution of 1.25×1.25×1.25 mm^3^. We used only the b = 2000 s/mm^2^.

#### Human diffusion data: STN96 dataset (subject H4)

This dataset was acquired at Stanford’s Center for Cognitive and Neurobiological Imaging (www.cni.stanford.edu). The dMRI data were sampled in 96 directions and the spatial resolution is 1.5×1.5×1.5 mm^3^. The b-value was set to 2000 s/mm^2^. FMRI measurement of the visual field maps (Dumoulin and Wandell 2008) and T1-weighted anatomical image were also acquired from the same individuals. Informed written consent was obtained from all subjects.

The experimental procedures were approved by the Stanford University Institutional Review Board. This dataset is used in the analysis of previous papers (STN96 dataset; Pestilli et al. 2014; Rokem et al. 2015; Takemura et al. 2016a, 2016b).

### MR data preprocessing

#### Macaque diffusion data 1

Macaque diffusion data1 was preprocessed by the TORTOISE software package (Pierpaoli et al. 2010) for eddy-current and motion corrections. Processing details are presented in a previous publication (Thomas et al. 2014).

#### Macaque diffusion data 2-4

Eddy-currents and motion correction was performed by a 14-parameter constrained nonlinear co-registration based on the expected pattern of eddy-current distortions given the phase-encoding direction of the acquired data (Rohde et al. 2004). The direction of the diffusion-gradient in each diffusion-weighted volume was corrected using the rotation parameters from the motion and eddy-current distortion correction procedure. All pre-processing steps have been implemented in Matlab as part of the mrVista software distribution (https://github.com/vistalab/vistasoft).

The dMRI data were registered to the mean of the (motion-corrected) non-diffusion-weighted (b0) images using a two-stage coarse-to-fine approach that maximized the normalized mutual information (Friston and Ashburner 2004). The mean non-diffusion weighted image was aligned automatically to the anatomical image using a rigid body mutual information algorithm.

#### Human STN96 data

For Human STN96 dataset, we also used the measurements of the B0 magnetic field for post-hoc correction of EPI spatial distortion (https://github.com/kendrickkay/preprocessfmri). Eddy-current correction was not applied because the data is corrected by using dual-spin echo sequence which minimizes eddy-current distortion (Reese et al. 2003). The other preprocessing step was identical to macaque diffusion data 2-4, and described in a previous publication (Takemura et al., 2016a).

#### Human HCP90 data

The Human Connectome data were preprocessed by the consortium using methods that are presented in a previous publication (Sotiropoulos et al. 2013).

### Tractography and fascicle evaluation

#### Connectome optimization

We used Ensemble Tractography (ET) to estimate the streamlines in the human and macaque data (Takemura et al. 2016a; https://www.github.com/brain-life/pestillilab_projects/et). This method begins by generating a large set of candidate streamlines using MRtrix (Tournier et al. 2012) using the entire white-matter volume as a seed region. We generated a candidate connectome using probabilistic tractography and five curvature thresholds (minimum radius of curvature, 0.25, 0.5, 1, 2, and 4 mm; Takemura et al. 2016a). We used Constrained Spherical Deconvolution (CSD; Tournier et al. 2007) for tractography and set the maximum number of harmonics in CSD to 8 (*L_max_*=8) for all datasets. We set other parameters as default (step size: 0.2 mm; maximum length: 200 mm; minimum length: 10 mm). We generated a total of 10,000,000 streamlines for the large human brain and 2,500,000 streamlines for the smaller macaque brain. Finally, we used Linear Fascicle Evaluation (LiFE; Pestilli et al. 2014; Caiafa and Pestilli, 2015) to optimize the candidate connectome. This procedure removes the streamlines that make no significant contribution to explaining the diffusion measurements. The number of streamlines retained in an optimized connectome depends on the quality and spatial resolution of the data, ranging from 45,969 (M4, 1 mm isotropic) to 925,593 (M1, 0.25 mm isotropic) streamlines in the macaque data, and from 266,246 (H4, STN96, 1.5 mm isotropic) to 450,966 (H2, HCP90, 1.25 mm isotropic) in the human data.

The data quality and spatial resolution in the M1 (post-mortem) data is extremely high. Even though the volume of the macaque brain is only 7% of the human brain, the spatial resolution of the M4 data is 125 times greater than the high resolution HCP human data. Hence, the number of streamlines retained by LiFE in the M1 dataset exceeds the number retained to model the human dataset.

### Cortical area identification

*Human data.* For STN96 dataset, we identified the location of cortical areas (visual field maps) using fMRI data and population receptive field method (Dumoulin and Wandell 2008; Wandell and Winawer 2011). We identified the border between visual areas (V1, V2, V3, hV4, V3A/B, VO, LO, TO and IPS-0) based on the reversal of polar angle, eccentricity and anatomical landmarks (Press et al. 2001; Dougherty et al. 2003; Brewer et al. 2005; Larsson and Heeger 2006; Amano et al. 2009; Witthoft et al. 2014; Winawer and Witthoft 2015). Technical details of the method are described in previous publications (Dumoulin and Wandell 2008; Amano et al. 2009; Winawer et al. 2010; Takemura et al. 2012).

For the HCP90 dataset, we identified the location of visual field maps based on the surface-based probabilistic atlas proposed by Wang and colleagues (Wang et al. 2015). We adapted the visual field maps defined in the atlas (V1/V2/V3, V3A/B, hV4, LO, TO and IPS-0) for the T1-weighted image in individual HCP90 dataset based on surface-based registration.

*Macaque data.* We used the Saleem and Logothetis (Saleem and Logothetis 2012) MR atlas to identify areas in macaque visual cortex (V1, V2, V3, V3A, V4d, V4v, MT and TEO). Briefly, we compared the coronal slice of the anatomical image (T1-weighted, T2-weighted or b=0 image) with the coronal section of the atlas comparing the area identification based on the histology and anatomical MRI data. This method is the same as a previous study (Thomas et al. 2014).

### Tract identification

For human data, we identified major white matter tracts in occipital cortex using Automatic Fiber Quantification (AFQ; Yeatman et al. 2012b; https://github.com/yeatmanlab/AFQ). AFQ defines waypoint ROIs in individual subject by non-linear transformation from waypoint ROI in MNI template brain, which is drawn on the basis of anatomical prescription (Wakana et al. 2004). The streamlines that pass through a pair of ROIs are considered as potential members of a given tract. The set of potential streamlines is refined by removing outliers. These are streamlines that meet the following criteria: (1) the streamline length ≥ 3 sd longer than the mean streamline length in the tract, (2) the streamline position is ≥ 3 sd away from the mean position of the tract (Yeatman et al. 2012b).

For macaque data, we used waypoint ROIs in structural images based on anatomical prescription to identify major tracts in human and macaque occipital cortex (Catani et al. 2002; Wakana et al. 2004).

There were a few additional processing steps. First, we used conTrack (Sherbondy et al. 2008a) to identify the human OR. Second, in both human and macaque VOF, we added a constraint that the streamlines must be dorsal-ventral between the two ROI waypoints. Streamlines whose path deviated more than 2 sd from the mean direction of the VOF streamlines (Takemura et al. 2016b) were deleted. Finally, tract visualization is implemented in the Matlab Brain Anatomy toolbox (https://github.com/francopestilli/mba; Pestilli et al., 2014).

#### Human data

*ILF, IFOF and Forceps Major.* ILF, IFOF and Forceps Major are defined as the tract passing through two waypoint ROIs derived from AFQ. For some subjects, we manually edited the position of waypoint ROI when MNI transformation produces ROIs in an erroneous position. We finally select streamlines in the optimized connectome passing through two waypoint ROIs. For the ILF, we also tested five different waypoint ROI definitions to examine the generality of the results (see Supplementary Figure 6).

*VOF.* We identified the VOF using the VOF toolbox within AFQ software package (https://github.com/jyeatman/AFQ/tree/master/vof). Briefly, this method identifies the location of ventral occipital-temporal cortex as a ROI using the Freesurfer (Fischl 2012; http://surfer.nmr.mgh.harvard.edu) parcellation, and then select streamlines having endpoint in the ROI from the optimized connectome. The detailed explanation of the VOF identification method is described in previous articles (Yeatman et al. 2014; Duan et al. 2015; Weiner et al., 2016a).

*OR.* We used conTrack (Sherbondy et al. 2008a) to estimate the human OR. First, we manually estimated the approximate location of the Lateral Geniculate Nucleus (LGN) on T1-weighted image. To confirm the manual definition of the LGN, we confirmed that streamlines originating in the optic chiasm terminate in the LGN ROIs by using deterministic tractography (Ogawa et al. 2014). We then placed a 5-mm radius sphere that covers the LGN endpoints of streamlines from the optic chiasm. Second, we identify the location of the V1 using Freesurfer (Hinds et al. 2008; Fischl 2012). We generated 100,000 candidate streamlines connecting between the LGN and V1 using conTrack, and selected 5,000 streamlines with the highest conTrack scores. Streamlines were only excluded if they traversed a non-biological path, such as passing through the ventricles or crossing to the other hemisphere. Further details on the methods to identify the OR using conTrack are described in previous papers (Sherbondy et al. 2008b; Levin et al. 2010; Ogawa et al. 2014; Duan et al. 2015).

#### Macaque data

For macaque, we manually defined the waypoint ROI based on the anatomical prescription reported in a previous work (Schmahmann and Pandya 2006).

*ILF.* We identified the macaque ILF using two-plane “AND” ROIs (anterior, posterior), which are manually drawn in the coronal plane on the basis of the previous study (Schmahmann and Pandya 2006). We also manually defined the “NOT” ROIs covering the Sagittal Stratum in the coronal plane, which is visible in high-resolution non-diffusion weighted image (b=0), following anatomical prescription (Schmahmann and Pandya 2006). The ILF is defined as a group of streamlines passing through two “AND” waypoint ILF ROIs, and does not pass through “NOT” ROIs defining the Sagittal Stratum. See Supplementary Figure 1A to see the location of waypoint ROIs.

*Forceps Major.* We identified the macaque Forceps Major using three waypoint ROIs. The first ROI is the corpus callosum which is defined manually from an anatomical image. The second and third ROIs are coronal planes in the left and right hemispheres; each plane is located near the anterior edge of MST as defined in the atlas (Saleem and Logothetis 2012). The Forceps Major is defined as a group of streamlines passing through all three waypoint ROIs.

*OR.* To identify the OR we manually identified the LGN from high-resolution non-diffusion weighted image (b=0) and V1 from MRI-based atlas (Saleem and Logothetis 2012). We identified the OR as a group of streamlines having one endpoint in the LGN and the other endpoint within 3 mm of the V1 gray matter ROI.

*VOF.* We identified the macaque VOF using two waypoint ROIs in axial plane (dorsal, ventral; Takemura et al. 2016b). Supplementary Figure 1B shows a waypoint ROI placed in the left hemisphere of M1. We placed dorsal plane inferior to the Angular Gyrus, because Wernicke (1881) indicated that the dorsal projection of mVOF may include the Angular Gyrus. The posterior limit of dorsal plane is the lunate sulcus, and the anterior limit is the superior temporal sulcus. The position of the ventral plane is dorsal to occipital temporal sulcus. The anterior limit of the ventral plane is also the STS, whereas the posterior limit is the inferior occipital sulcus.

The macaque VOF is the group of streamlines passing through both planes (see Supplementary Figure 1A for the location of waypoint ROIs).

#### Virtual lesions

There are many possible ways that one could assess the statistical evidence supporting a collection of streamlines. Here, we used the Virtual Lesion method (Pestilli et al. 2014; Takemura et al. 2016b; Leong et al. 2016) to evaluate the evidence supporting the existence of the macaque VOF. Specifically, we compare the change in prediction accuracy (root mean squared error; RMSE) for diffusion signal between the optimized connectome and a lesioned connectome with the streamlines of interest removed. The RMSE is compared in all voxels touched by the lesioned streamlines, the mVOF in this case. The complete set of streamlines that contribute to the prediction of the diffusion measurements in these voxels is called the path-neighborhood of the mVOF. This path neighborhood includes the mVOF itself and all of the other streamlines that pass through the mVOF voxels. We measure the distribution of RMSE values in the mVOF voxels when using the entire path-neighborhood and then we remove the mVOF and solve for the weights with the remaining streamlines. The supporting evidence is assessed with two different measures. We compare the two RMSE distributions using the Earth Mover’s Distance (*EM*), and also by calculating the strength of evidence, *S,* which is the difference in the mean RMSEs divided by the joint standard deviation (see Pestilli et al. 2014).

#### Relating the VOF endpoint and cortical area

Streamlines terminate at the boundary between white and gray matter. We measured the distance between tract endpoints and gray matter voxels. We could then identify the cortical areas closest to the tract endpoints. Specifically, we collected the X, Y, and Z coordinates of the endpoints of the VOF streamlines and computed the distance between the endpoints and gray matter voxels. For each gray matter voxel, we counted the number of endpoints within a threshold distance (human data: 3 mm, macaque data: 2 mm). We plot the normalized endpoint counts on the smoothed cortical surface (Figure 6).

This analysis has some limitations, derived from the challenges in associating the cortical surface and tract endpoints (Reveley et al. 2015). This analysis measures only the general proximity between cortical maps and tract endpoints, rather than the definitive estimates of the fiber projections into cortical gray matter surface.

## Results

We begin by comparing the spatial arrangement of the major white matter tracts in both human and macaque visual cortex. We focus much of our attention on a tract that has been little studied: the human and macaque VOF. We then identify its position with respect to other tracts and the location of its endpoints with respect to the cortical maps.

### Comparison of major occipital tract positions

Figure 2 compares the general organization of the major occipital tracts in macaques (Figure 2A) and humans (Figure 2B). We consistently identified the major tracts in both datasets (OR, ILF, Forceps Major and VOF) from both datasets. These tracts are reported in previous studies in human (Wakana et al. 2004; Catani and Thiebaut de Schotten 2008, 2012; Martino and Garcia-Porrero 2013; Yeatman et al. 2013; Takemura et al. 2016b) and macaque (Schmahmann and Pandya 2006; Schmahmann et al. 2007), with the exception of the macaque VOF. We discuss this tract in more detail in the next section.

**Figure 1.**
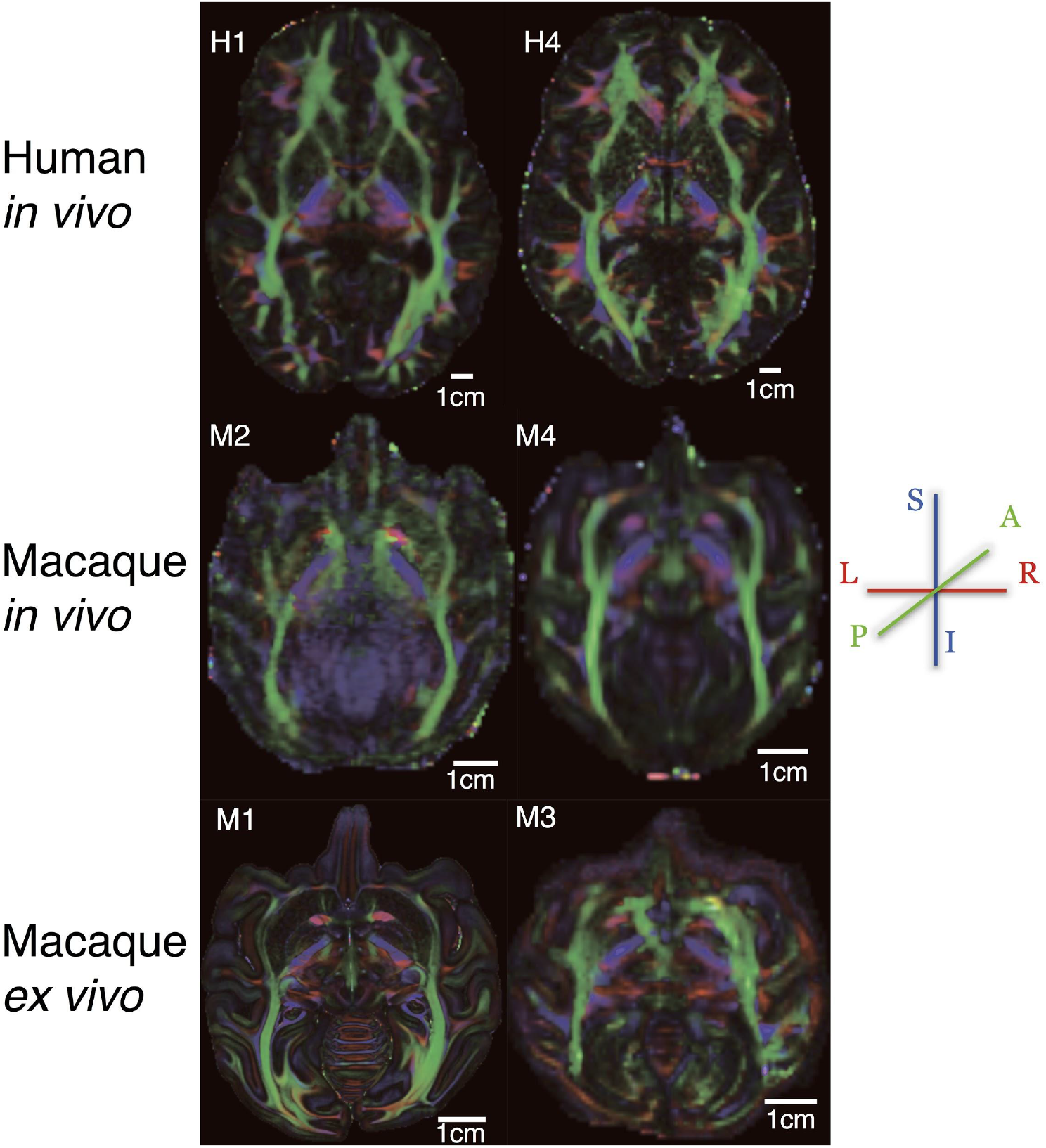
The resolution of the several diffusion MRI data sets. Panels show axial-plane images of the Principal Diffusion Direction (PDD) map of six representative subjects in human and macaque datasets. Data were acquired at different angular and spatial resolution (see Materials and Methods). Colors indicate the PDD in individual voxels (blue: superior-inferior, S-I; green: anterior-posterior, A-P; red: left-right, L-R). Top panel shows the PDD in two living human datasets; one in HCP90 (H1, 1.25 mm isotropic) and the other in STN96 dataset (H4, 1.5 mm isotropic) dataset. The middle panels are the datasets from living macaque (left: M2, 0.75 mm; right: M4, 1 mm). The bottom panels are datasets from post-mortem macaque (left: M1, 0.25 mm; right: M3, 0.8 mm). In later section, we describe the dependency of results on spatial resolution among datasets (Figures 8 and 9).

**Figure 2.**
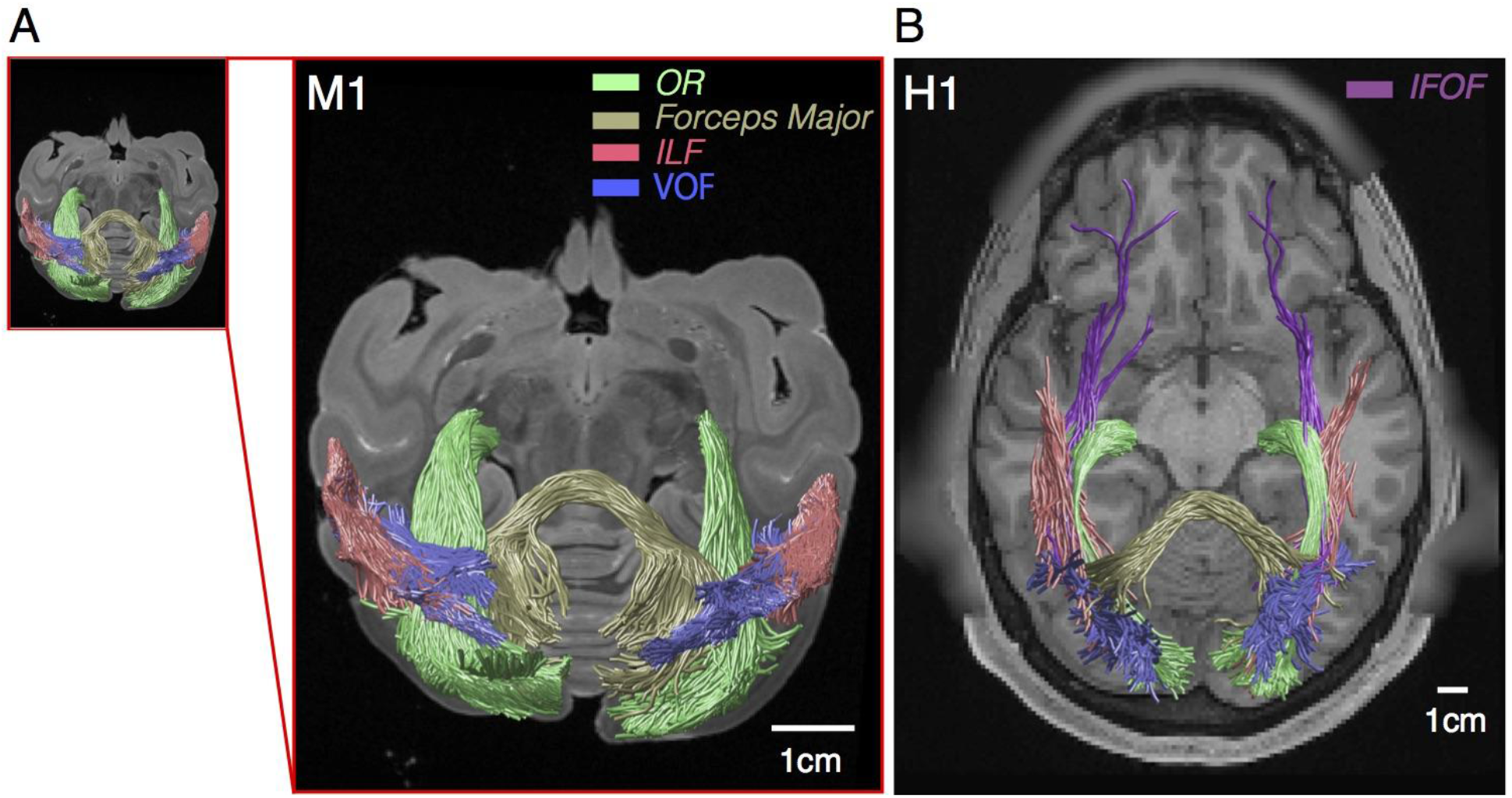
Major white matter tracts with at least one endpoint in occipital cortex. Axial view of major white matter tracts (ILF, red; IFOF, purple; OR, green; Forceps Major, dark yellow; VOF, blue) overlaid on anatomical image in a representative macaque (**A,** subject M1, *ex vivo*) and human (**B**, subject H1, HCP90 dataset). The small panel in the left side in panel A indicates the size of the macaque tracts in an identical spatial scaling with human figure.

There is a significant difference between the major occipital tracts in human and macaque: the macaque dMRI data does not provide a strong support for the IFOF. This is consistent with previous studies showing evidence for the existence of the IFOF in humans (Catani and Thiebaut de Schotten 2008; Martino et al. 2010a; 2010b; 2011; Sarubbo et al. 2013; Pestilli et al. 2014; Forkel et al. 2014), but no evidence for this tract in macaques (Schmahmann and Pandya 2006; Schmahmann et al. 2007; but see Mars et al. 2015).

### Identification of macaque VOF (mVOF)

Figure 3 shows the Principal Diffusion Direction (PDD) map in the highest resolution *ex vivo* macaque dMRI dataset (subject M1, 0.25 mm isotropic voxels; see Supplementary Figure 2 for other slices). PDD map has been used to identify the position of the tract in human studies (Pajevic and Pierpaoli 1999; Wakana et al. 2004; Yeatman et al. 2013; Takemura et al. 2016b). Inspection of the PDD images clearly reveals the position of several major tracts; such as the OR and Forceps Major (Figure 3 and Supplementary Figure 2).

**Figure 3.**
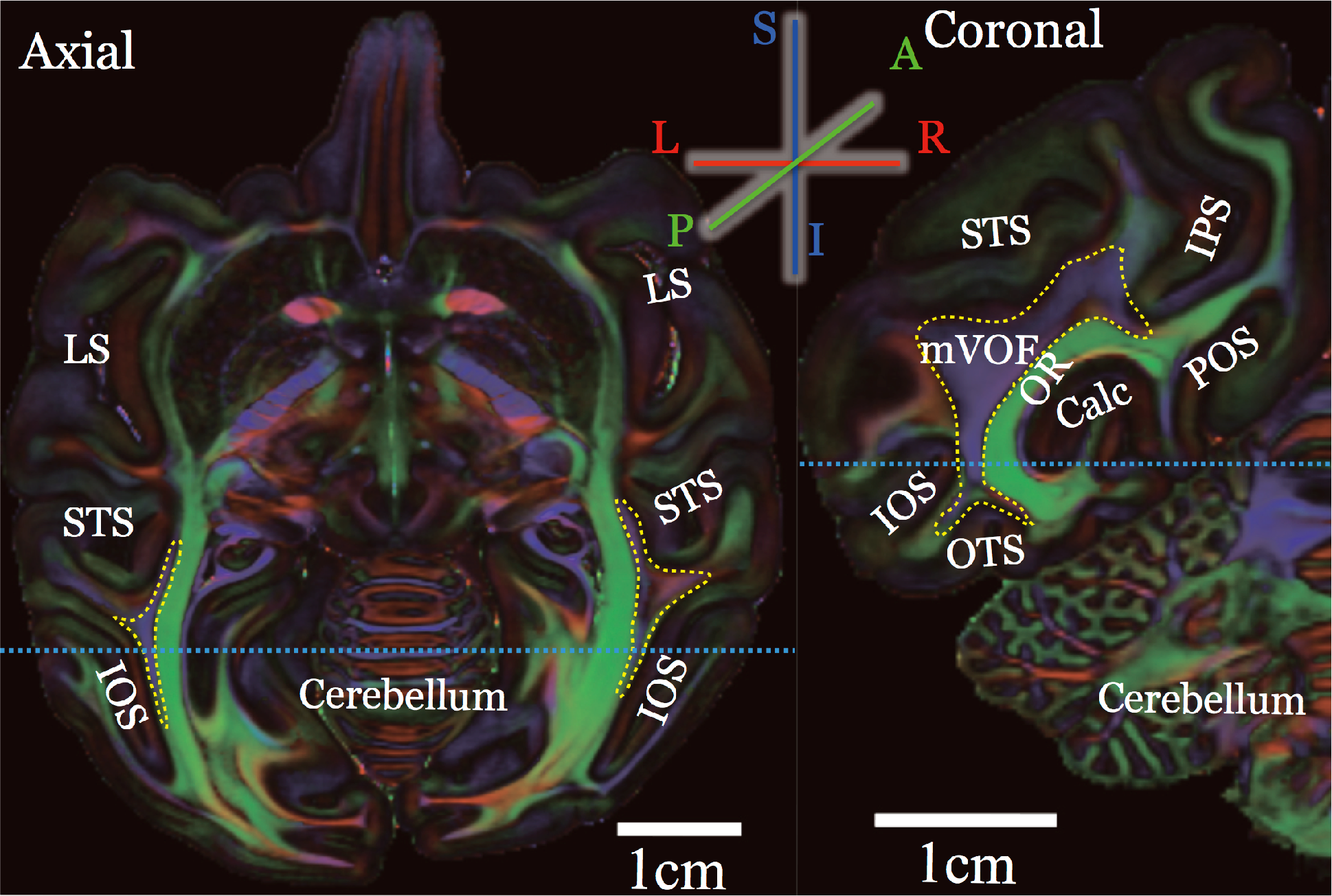
The position of the macaque VOF. The position of the mVOF identified in PDD map (subject M1, *ex vivo*). The color scheme depicts the PDD in each voxel (blue, superior-inferior; green, anterior-posterior; red, left-right). In the axial slice (left panel), we could see the white matter portion with predominantly superior-inferior diffusion signal between Superior Temporal Sulcus (STS) and Inferior Occipital Sulcus (IOS). In the coronal slice (right panel), this region is located lateral to the Calcarine sulcus and the Optic Radiation (OR, which has anterior-posterior PDD, green). Scale bar (white) in each panel indicates 10 mm. Light blue dotted line in left panel indicates the position of coronal slice in the right panel, vice versa. LS: Lateral Sulcus, IPS: Intraparietal sulcus, Calc: Calcarine sulcus, POS: Parieto-Occipital Sulcus, OTS: Occipito-Temporal Sulcus.

Importantly, the PDD maps in this high resolution dataset clearly identify the major macaque pathways including a pathway seemingly homologous to the human VOF. Specifically, the data in Figure 3 reveal tracts with a superior-inferior diffusion direction (blue) in the lateral occipital white matter (outlined). The blue PDD indicates the presence of a vertical tract communicating between dorsal and ventral visual cortex. The ventral portion of this tract is located between Inferior Occipital Sulcus (IOS) and the Superior Temporal Sulcus (STS; left panel, axial view, Figure 3). This tract is lateral to the Calcarine sulcus and the OR (green regions in right panel, Figure 3). This tract is also identifiable from PDD map in other slices (Supplementary Figure 2).

Hereafter, we call this tract macaque Vertical Occipital Fasciculus or mVOF. We consistently identified the core portion of this tract from the PDD map in the dataset from living and post-mortem macaque brain with coarser spatial resolution (see Figure 8; “Instrumentation and acquisition parameter dependencies”). We also note that the position of mVOF is qualitatively consistent with classical studies by Wernicke and other authors (Wernicke 1881; Bailey et al. 1944; Petr et al. 1949; see “Diffusion MRI estimates of the mVOF are consistent with anatomical studies” in Discussion).

Figure 4A describes the estimated mVOF visualized as a tract using tractography (see Materials and Methods). The mVOF is located on the anterior-lateral side of the lunate sulcus dorsally and its ventral endpoints are near the occipito-temporal sulcus (OTS). We observed the core of the mVOF in this position for all of the macaque brains (see Figure 8 for more examples). Figure 4B is the human VOF identified from HCP90 dataset, shown for a comparison.

**Figure 4.**
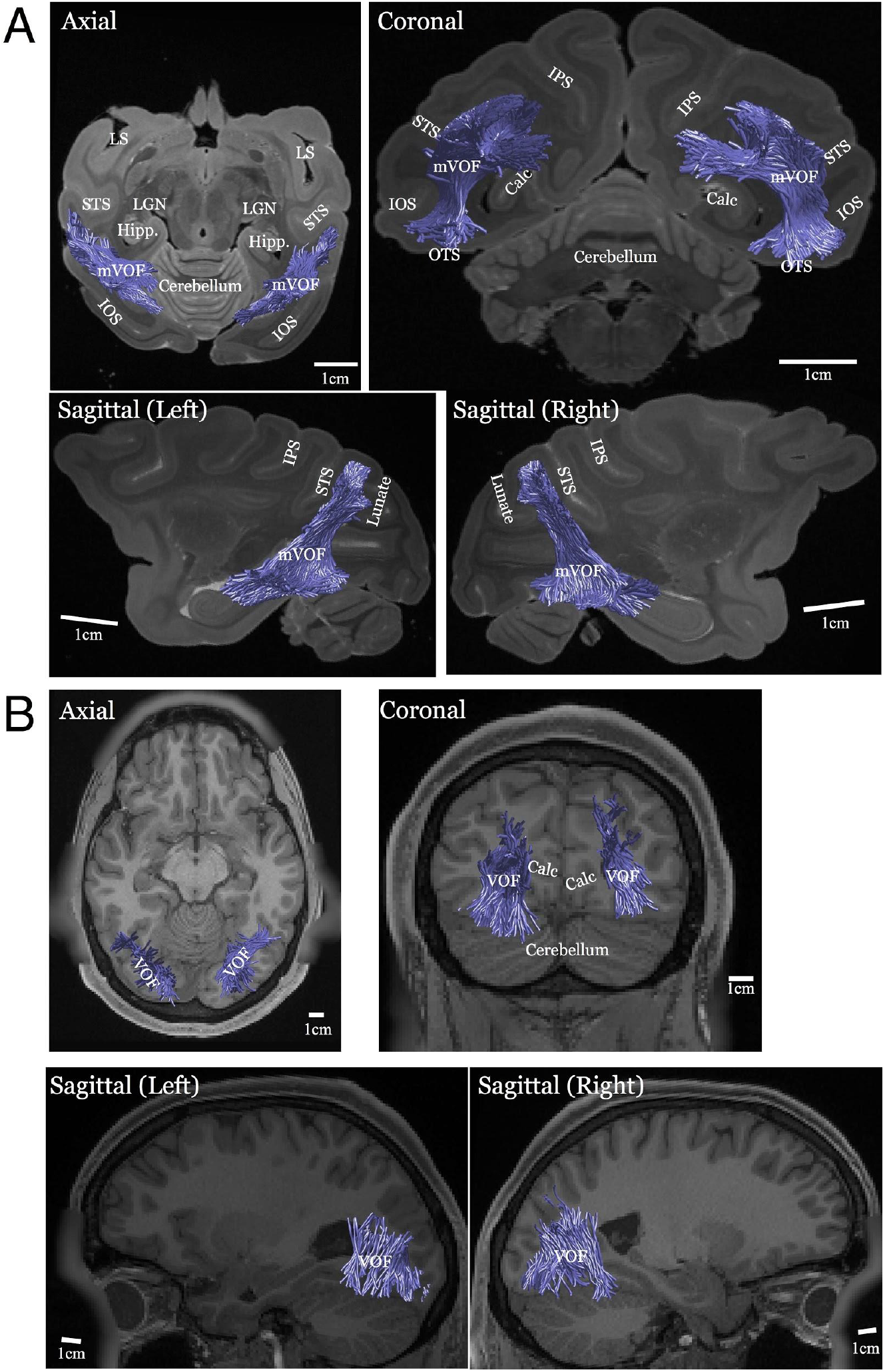
Human and macaque VOF identified using tractography. **A.** Macaque VOF identified using tractography, overlaid on non-diffusion weighted (b=0) image (subject M1; left top, axial slice; right top, coronal slice; bottom, sagittal slices). Lunate, Lunate Sulcus, Hipp: Hippocampus. Other conventions are identical to those in Figure 2. **B.** Human VOF identified using tractography, overlaid on T1-weighted image (subject H1, HCP90dataset). Scale bar (white line) in each panel indicate 10 mm.

#### Statistical evidence in support of the mVOF tract

We use the virtual lesion method (Honey and Sporns 2008; Pestilli et al. 2014; Takemura et al. 2016b; Leong et al. 2016) to evaluate the strength of evidence supporting the existence of the mVOF. Briefly, the virtual lesion method compares the prediction accuracy on the diffusion signals between two connectome models. The first connectome model contains all streamlines (unlesioned), and the second connectome model contains all streamlines except the tract of interest (lesion). First, we identify all voxels touched by the tract of interest (right mVOF, 1919 streamlines). We then find all of the rest of the streamlines passing through these voxels (path-neighborhood, 32399 streamlines). Finally, we compare the ability of the two models to predict the diffusion data within the mVOF voxels. More specifically, we compare the root mean square error (RMSE) in the mVOF voxels of the lesioned and unlesioned connectome models.

We compare the distributions of RMSE for lesioned and unlesioned models. First, we plot the distribution of root mean squared error (RMSE) between these two models in subject M1; the model with and without the right mVOF (Figure 5). There is a clear difference in the RMSE distribution of the two models, with the lesioned model doing a substantially worse job. We quantify the difference between these distributions by computing the strength of evidence (*S*) and Earth Mover’s distance (*EM*) as an index for statistical support for this pathway (Pestilli et al. 2014). Figure 5 shows that there is a high strength of evidence for the mVOF pathway in the right hemisphere (S = 14.03). There is also a large earth mover’s distance between the two RMSE distributions (EM = 4.24) for right mVOF in subject M1. The strength of evidence is also significant in the left mVOF (S = 25.63; EM=6.3). Thus, in addition to visible evidence on PDD maps (Figure 3 and Supplementary Figure 2), there is a strong statistical support for the mVOF.

**Figures 5.**
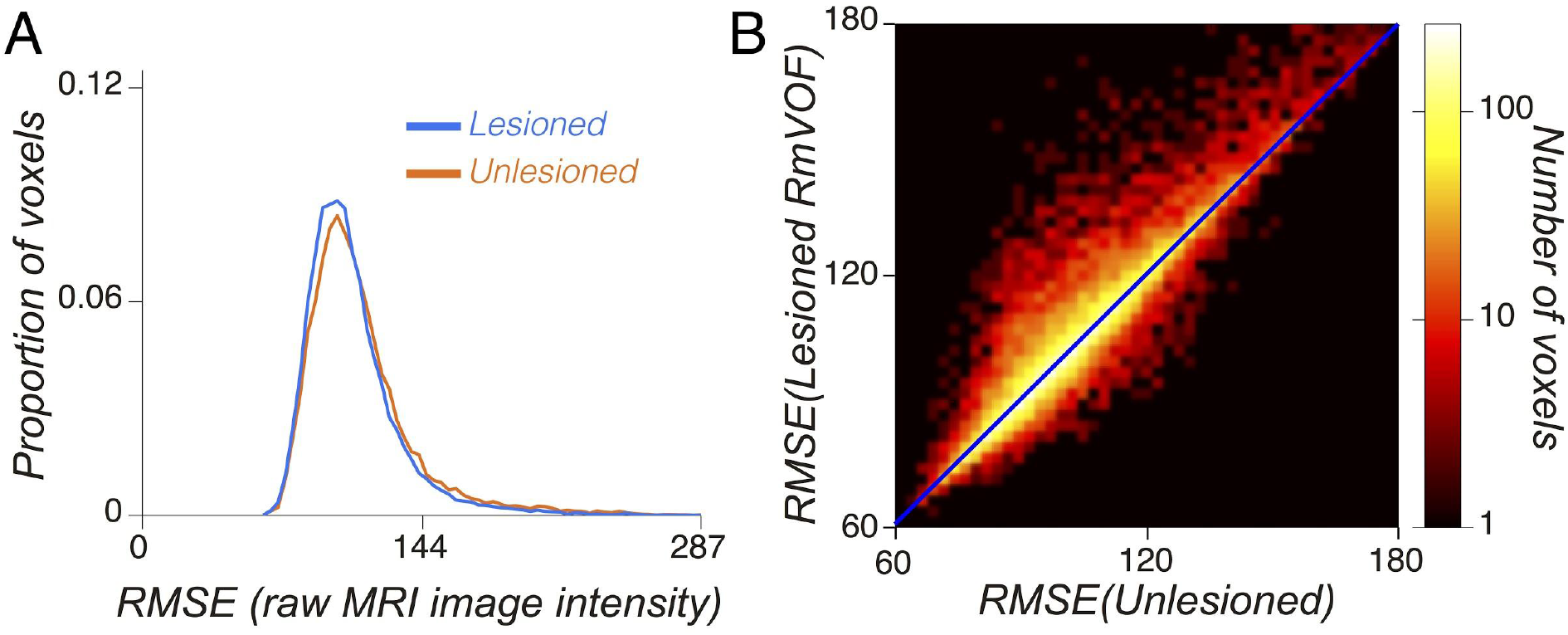
Statistical evidence supporting the existence of the mVOF. **A.** The histogram of the model accuracy (root mean squared error; RMSE) of two models (red; unlesioned model, blue; lesioned model) in voxels along the right mVOF or its path-neighborhood, in subject M1 (see Material and Methods). The difference of two distributions is statistically significant (*S* = 14.03, *EM* = 4.24). **B.** Model accuracy comparison between unlesioned and lesioned models in voxels along right mVOF in subject M1 (abscissa: unlesioned model, ordinate: lesioned model). Color map indicates number of voxels.

#### Cortical regions near the mVOF endpoints

Next, we identified the cortical maps near the endpoints of the VOF in humans and macaques. In humans, visual field maps were identified using fMRI as reported in a previous study (Takemura et al. 2016b) as well as using a surface-based atlas (Wang et al. 2015). In macaques, visual areas were identified using a standard MRI-based atlas (Saleem and Logothetis 2012). Tract cortical endpoints were computed by counting the number of VOF streamlines having endpoint near each gray matter voxel (tract endpoint density, see Materials and Methods; Takemura et al. 2016b). Normalized streamline endpoint density was superimposed on the cortical surface (Figure 6; see Methods).

**Figure 6.**
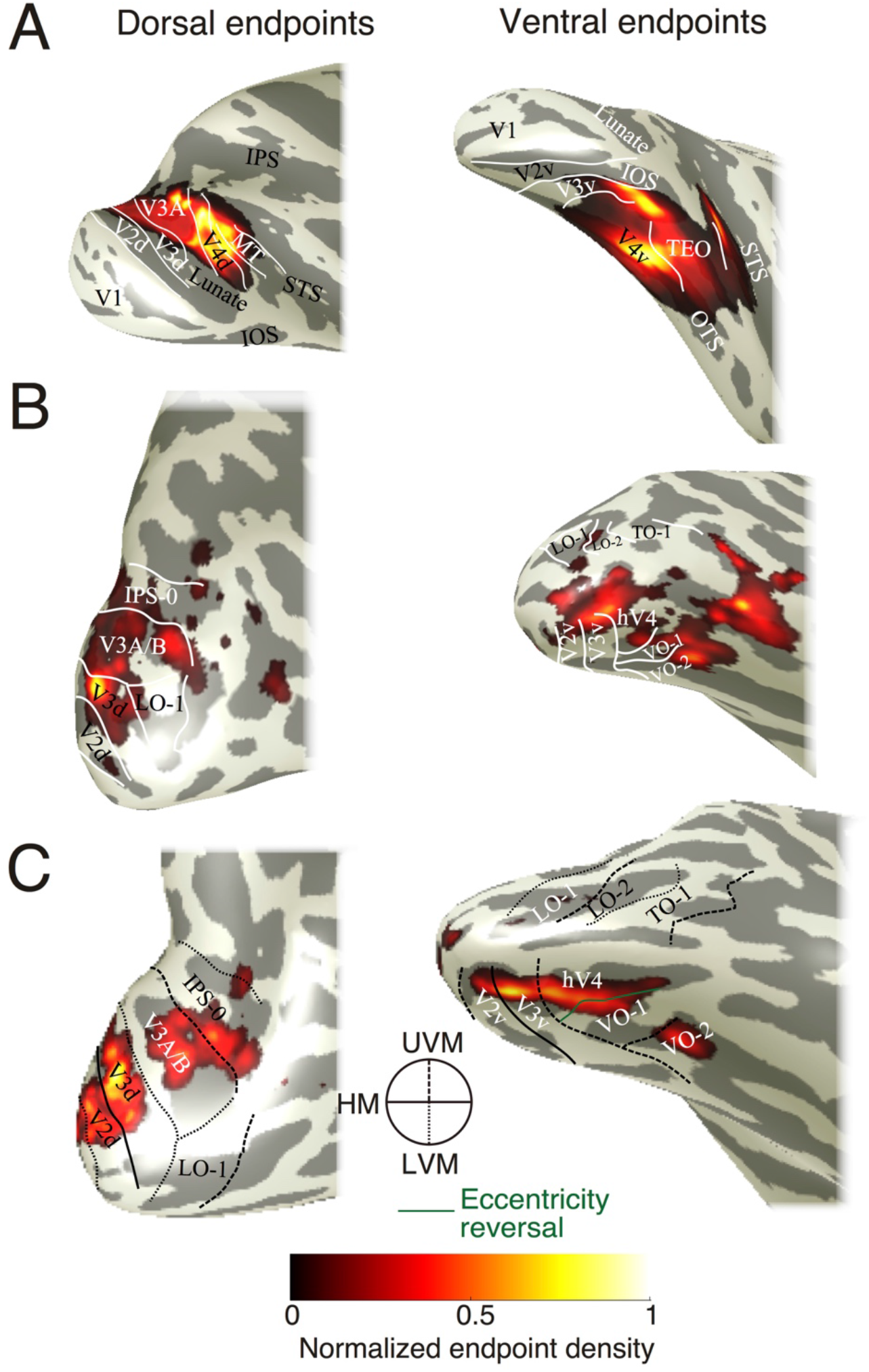
Cortical visual maps near VOF endpoints in human and macaque. **A.** Cortical maps near the mVOF endpoint (subject M1, right hemisphere). In this plot, we calculated the spatial distance between mVOF endpoints and gray matter voxels, and counted the number of mVOF streamlines having endpoints close to the gray matter voxel (see Materials and Methods). The boundaries between visual areas are manually identified using a MRI-based atlas (Saleem and Logothetis 2012). We observed the putative mVOF termination near V3A, V4d dorsally and V4v, TEO ventrally. See Figure 9 and Supplementary Figure 3 for more examples. **B.** Cortical maps near human VOF endpoint in HCP90 dataset, Subject H1). The boundaries between cortical areas are estimated using a surface-based probabilistic atlas (Wang et al. 2015). **C.** Cortical maps near human VOF endpoint in relation to visual field maps estimated individually using fMRI (Dumoulin and Wandell 2008; Wandell and Winawer 2011; STN96 dataset, Subject H4). The boundaries between cortical areas are defined by the boundaries of polar angle and eccentricity estimated by fMRI-based visual field mapping (see captions at the center; circular inset; UVM, upper vertical meridian; HM, horizontal meridian; LVM, lower vertical meridian).

Figure 6A shows the mVOF terminations in several visual field maps (right hemisphere, subject M1). One end of the mVOF terminates near the anterior portion of the Lunate sulcus dorsally, and OTS ventrally. The left panel of Figure 6A shows maps near dorsal mVOF endpoints. The region includes V3A, V4d/DP and possibly MT (see Figure 8 for a comparison of the consistency across datasets, and Supplementary Figure 3 for the results in left hemisphere). Right panel of the Figure 6A shows the maps near ventral mVOF endpoints. Endpoints are near cortical regions around OTS, which is mostly V4v. There are some endpoints near the anterior portion of the OTS, which may correspond to TEO (Saleem and Logothetis 2012; Kolster et al. 2014; see Supplementary Fig 3 for the results in left hemisphere). We note that the estimates of cortical endpoints depend on the spatial resolution of the diffusion MRI data (see “Instrumentation and acquisition parameter dependencies”; Figure 8).

Figures 6B and 6C describe the results on the right hemisphere in human dataset in two representative participants (subject H1, HCP90 dataset; Subject H4, STN96 dataset). The datasets are the same used previously (Takemura et al. 2016b), but analyses differed from previous reports because of the use of Ensemble Tractography methods (Takemura et al. 2016a; see Materials and Methods). New methods reproduce previous results (Takemura et al. 2016b), showing that the dorsal human VOF endpoints are proximal to V3A and V3B, V3d and IPS-0 whereas the ventral endpoints are proximal to human V4, VO and neighboring maps. This pattern is consistent across datasets (HCP90, Figure 6B and STN96, Figure 6C; see Supplementary Figure 4 for more examples). Human VOF endpoints supported by HCP90 dataset cover lager portions of cortical maps presumably because of the higher spatial resolution.

Whereas there are differences in visual field map organization between humans and macaques, results seem consistent between human and macaque VOF; the VOF endpoints are near V3A, and ventral endpoints are near ventral V4. These similarities in cortical endpoints suggest that the mVOF is homologous to human VOF.

#### VOF position with respect to other tracts

The human brain volume is about 15 times larger than the macaque brain, and the human cortical surface area is greatly expanded as well (Rilling 2006; Wandell and Winawer 2011; Corthout 2014). The structures in the human brain are not a one-to-one match with the macaque, simply scaled up (Rilling 2006; Passingham 2009). Investigators from many previous studies sought to associate the findings in macaque to the human brain and argued about the homology in functional measurements (Brewer et al. 2002; Fize et al. 2003; Tsao et al. 2003, 2008; Orban et al. 2004; Kriegeskorte et al. 2008; Wade et al. 2008; Kornblith et al. 2013; Kolster et al. 2014). Hereafter, we compare the gross anatomy on the human and macaque VOF.

Figure 7A shows PDD map in human (HCP90 H1) and macaque (M1). For ease of comparison, we used the same scale for both human and macaque PDD maps. We report noticeable difference between human and macaque VOF. The volume of human VOF is substantially larger than that of mVOF. The volume of the human brain is 15-times that of the macaque brain (Corthout 2014). Using the data from HCP90 dataset and M1, we estimate that the volume of the human VOF is approximately 31-times the volume of right mVOF. The relative increase in size of the human VOF seems to follow the general increase in area of the mid-level visual areas. We note that in macaque, early visual areas (such as V1) comprise a larger portion of the occipital cortex, whereas in humans mid-level visual maps are relatively larger (Brewer et al. 2002; Dougherty et al. 2003; Lyon and Connolly 2012).

**Figure 7.**
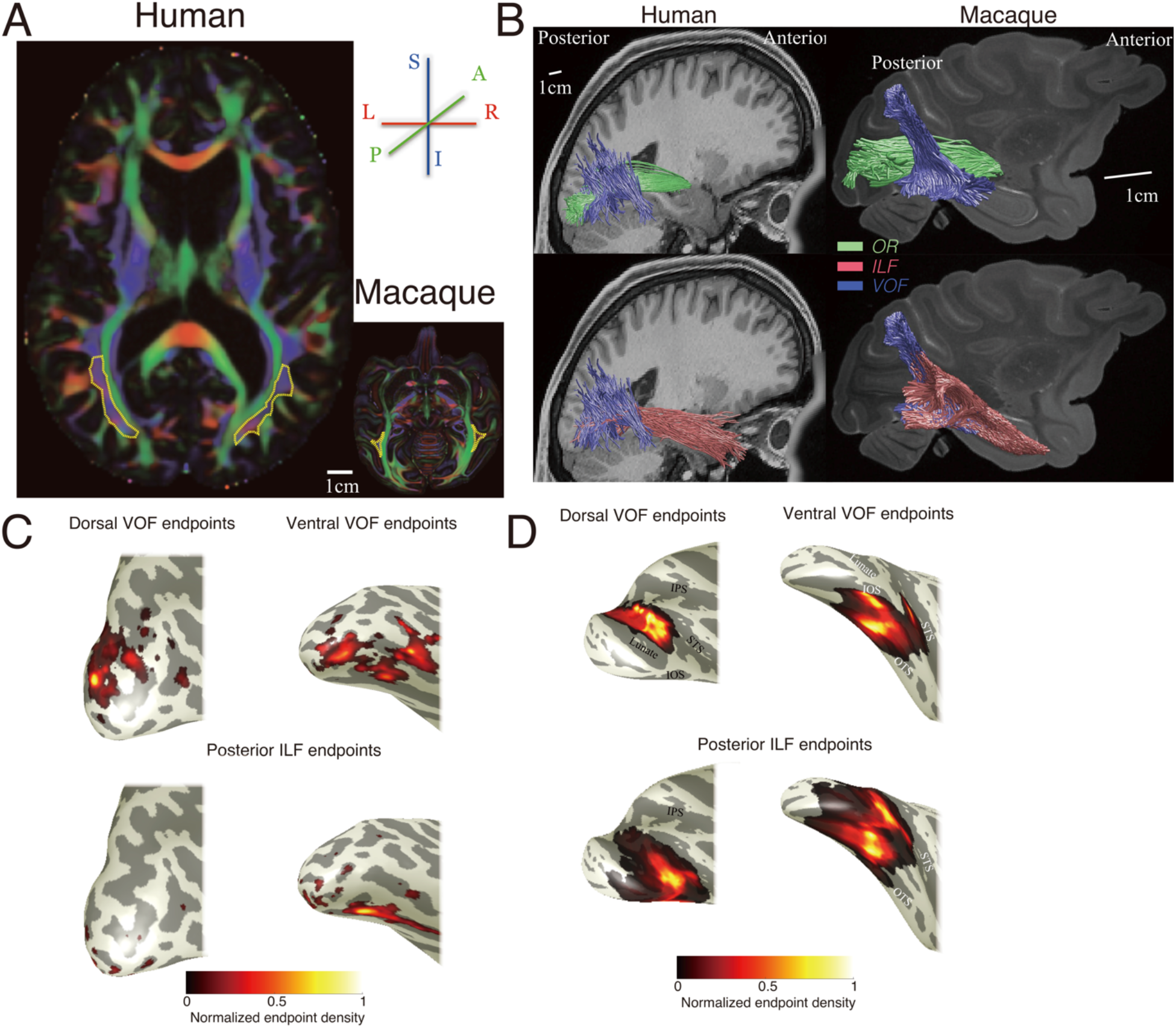
Human-macaque comparison. **A.** The comparison of the PDD map in human (subject H1, HCP90 data) and macaque (Subject M1) in an axial slice. Color schemes are identical to those in Figures 1 and 3. We plot the approximate position of the VOF by dotted yellow lines. While human VOF is elongated from anterior to posterior, the macaque VOF position is restricted between the IOS and OTS (see Figure 3 for details in macaque data). **B.** Spatial relationship between the Optic Radiation (OR; green), the Inferior Longitudinal Fasciculus (ILF, magenta) and VOF (blue) in human and macaque. **C-D.** VOF and ILF cortical endpoints in human and macaque (**C.** subject H1 from HCP90 dataset, **D.** subject M1).

Figure 7B plots the human and macaque VOF together with other fiber systems, the OR and the ILF. Human and macaque brains have similar relative position of VOF and OR. In both species, the VOF is located lateral to OR and they are clearly distinguishable (Figure 7B, top panels). This observation has been also reported in classical studies on post-mortem human and macaque (Wernicke 1881; Déjerine 1895).

We identified the macaque ILF using the guidelines in an earlier macaque dMRI and tracer studies (Schmahmann et al., 2007; Schmahmann and Pandya 2006). We then superimposed the VOF and ILF streamlines to investigate the relative position of these tracts.

We find that the relative position between ILF and VOF streamlines differs between human and macaque. A major portion of the macaque VOF communicates vertically between dorsal and ventral visual cortex (Figure 7B, right panel), and the dorsal end of the mVOF intermingles with the dorsal segment of the macaque ILF. For this reason, the mVOF cannot be well separated from the ILF (Figure 7B, bottom right). In human, the VOF is located lateral and orthogonally to the ILF streamlines. Most of the human VOF ventral streamlines can be distinguished from the ILF (Figure 7B, bottom left). There are a small number of intermingled streamlines in the human despite the fact that the human VOF is relatively large compared to the mVOF (Figure 7B; see Supplementary Figure 5 in other example from STN96 dataset). While estimated crossing between VOF and ILF depends on the selection for waypoint ROIs, however, we did not identify a large number of ILF streamlines intermingled with dorsal or ventral endpoint of the VOF streamlines (Supplementary Figure 6), unlike the results in macaque (Figure 7B).

Panels 7C and 7D show the positions of the dorsal and ventral VOF endpoints, as well as the posterior ILF endpoints, on the cortical surface. In this comparison, the estimated VOF and ILF endpoints are intermingled in the macaque while it is relatively distinguishable in the human.

### Instrumentation and acquisition parameter dependencies

The ability to resolve different tracts, estimate their sizes and endpoints depends strongly on the spatial and angular resolution of the acquisition (Kim et al. 2006; Roebroeck et al. 2008; Berman et al. 2013; Sotiropoulos et al. 2013, 2016; Pestilli et al. 2014; see also Lebel et al. 2012). The quality of the post-mortem data from subject M1 (Thomas et al. 2014; Reveley et al. 2015) is far beyond the other datasets. However, data collected from living brains offers additional opportunities even if it is limited in resolution (see “Combining post-mortem anatomical data with *in vivo* diffusion MRI” in Discussion). Here we tested what we can identify in relatively lower resolution datasets from macaque brains.

Figure 8A depicts the PDD map from macaque brain in other dMRI datasets. We can identify the mVOF (blue, superior-inferior) and the OR (Figure 8A; see Supplementary Figure 7 for a comparison in other slice selections). The result suggests that the core portion of the mVOF can be consistently identified in all macaque datasets, including *in vivo* data.

**Figure 8.**
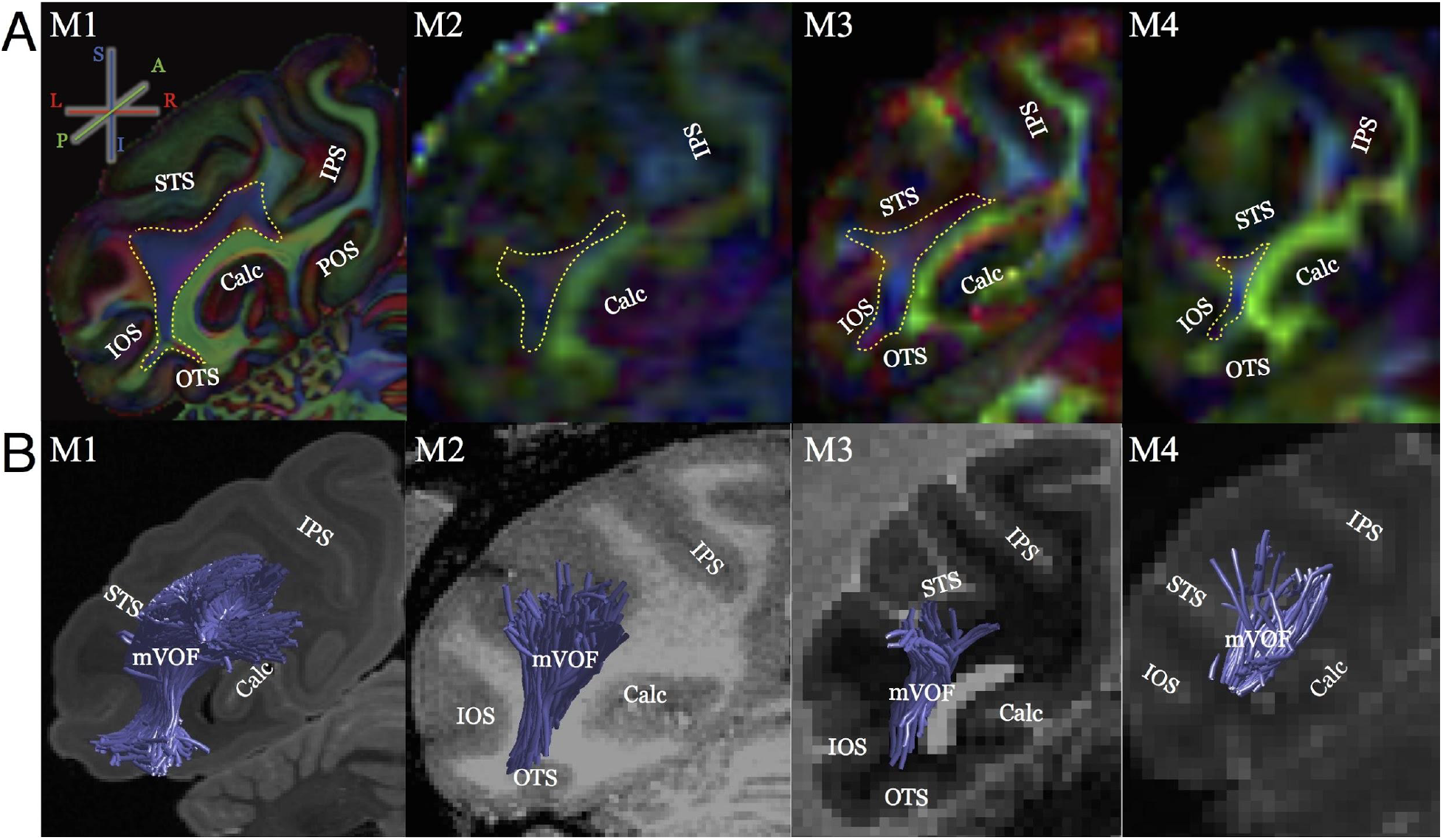
Consistency and dependency across the dataset with different resolutions. **A.** The comparison of left mVOF in PDD map across datasets. The color chart is identical to those used in Figure 3. **B.** The comparison of left mVOF identified by tractography across datasets.

While the datasets are qualitatively consistent, the estimates of the mVOF volume differ across datasets. For example, the mVOF volume estimated in subject M4 (*in vivo,* 1 mm isotropic, Figure 8B) is smaller than that in subject M1 (*ex vivo*, 0.25 isotropic, Figure 4A; Figure 8B) presumably because of the partial volume effect in lower resolution data. Depending on the data resolution, portions of even relatively large tracts can be missed.

The estimates of cortical endpoints also depend on spatial resolution. In subject M1 (0.25 mm isotropic; see also Figure 4) the mVOF branches enter a gyrus around the OTS; this branch is absent in the lower resolution data (Figure 8A). In subject M4 (1 mm isotropic), we could not identify mVOF endpoints close to the OTS (Figure 8A). These differences are probably not due to individual differences but to data resolution.

Figure 9 compares the cortical maps near mVOF endpoints among the datasets. While the center of the mVOF endpoints is consistent, mVOF endpoints in subject M1 (higher-resolution data) span a larger spatial extent on the cortical surface. In lower-resolution data, we particularly miss any cortical endpoints in the gyrus between IOS and OTS. This is because lower spatial resolution data is vulnerable to the partial volume effect with gray matter and superficial U-fiber system (Reveley et al. 2015; Vu et al. 2015). The results suggest that the improvement of spatial resolution on diffusion MRI data reduces the tractography bias for sulci over gyri; we still may miss some of the cortical endpoints in the dataset even with the highest spatial resolution used in the present study (Reveley et al. 2015; Sotiropoulos et al. 2016).

**Figure 9.**
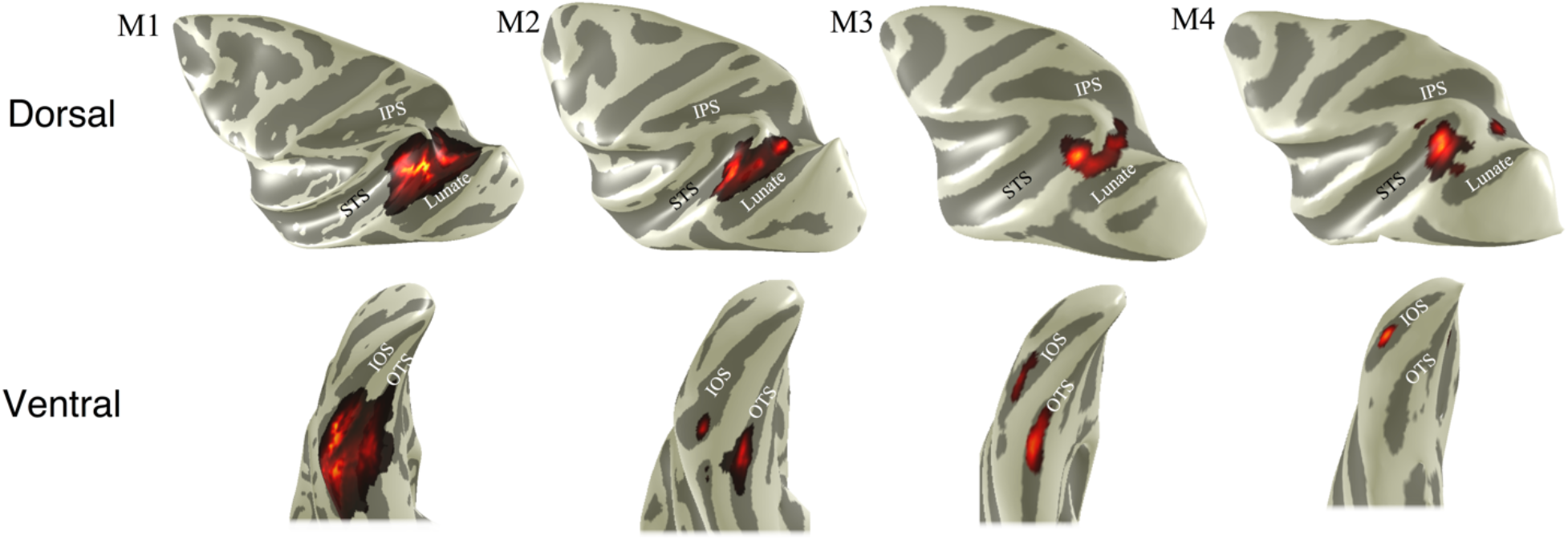
Spatial resolution dependency on the cortical endpoints estimate of mVOF. The comparison of estimated cortical mVOF endpoints across four macaque dMRI datasets collected at different spatial resolutions. The color map indicates the normalized density of left mVOF endpoint, as shown in Figure 6 (upper panels, dorsal mVOF endpoint; lower panels, ventral mVOF endpoint).

## Discussion

We used diffusion MRI data and tractography, in both human and macaque, to identify the major white matter tracts with at least one occipital endpoint. In both species we find apparently homologous tracts including the OR, Forceps Major, ILF and VOF. We find no evidence for a macaque IFOF, consistent with previous reports (Schmahmann et al. 2007; Catani and Thiebaut de Schotten 2008).

We succeeded in identifying the macaque VOF, which has been little studied. The core portion of this tract is consistent across dMRI datasets obtained with different spatial resolutions and number of diffusion directions. The macaque VOF shares some similarities with human VOF, particularly with regards to the position of its endpoints with respect to the cortical maps. But the position of the VOF in relation to the ILF may differ between species.

### Many homologous tracts, but no homologous IFOF

The dMRI analysis supports the existence of several large homologous tracts in macaque and human in occipital cortex: the OR, Forceps Major and ILF. This result supports the macaque dMRI analysis by Schmahmann and colleagues who also demonstrated consistency between the dMRI data and tracer study (Schmahmann et al. 2007).

There is also a notable difference between human and macaque occipital white matter tracts: the macaque dMRI data lack support for the IFOF, while the human dMRI supports the existence of an IFOF. This too is consistent with tracer (Schmahmann and Pandya 2006) and dMRI studies (Schmahmann et al. 2007) in macaque. The species difference concerning the IFOF has also been supported by classical and recent fiber dissection studies (Curran 1909; Martino et al. 2010a; 2010b; 2011; Sarubbo et al. 2013; Forkel et al. 2014). Further, the statistical support for the IFOF from dMRI data is highly significant (Pestilli et al. 2014). It remains uncertain whether human IFOF is composed by fully monosynaptic connections or series of connections (Mars et al. 2015). However that issue may resolve, there is a significant inter-species difference with respect to the estimated IFOF streamlines.

The functional significance of IFOF inter-species difference is an open and interesting question. One working hypothesis is that the IFOF is crucial for the rapid transmission of visual information to a semantic processing system in frontal cortex (Duffau et al. 2013). But the much greater size of the human brain, which is 15 times the volume of macaque, allows for many new functions and the possibility that the IFOF provides visual information to many of these circuits.

### Diffusion MRI estimates of the mVOF are consistent with anatomical studies

Wernicke (1881) reported the existence of a tract (*senkrechtes Occipitalbündel (fp)*; “perpendicular fasciculus”) connecting dorsal and ventral occipital cortex. His schematic of an axial slice of the post-mortem monkey brain is compared with the dMRI estimates of the principal diffusion direction (PDD) in macaque (Figure 10, “*fp*” in the right panel). Wernicke (1881) described the position of “perpendicular fasciculus” as lateral to the OR, which is consistent with PDD map (Figure 3). In Wernicke’s schematic diagram, the position of the perpendicular fasciculus is surrounded by two sulci, which correspond to modern definitions of the STS and the IOS. Thus, the existence and position of the mVOF derived from dMRI and tractography is consistent with Wernicke’s classical post-mortem study.

**Figure 10.**
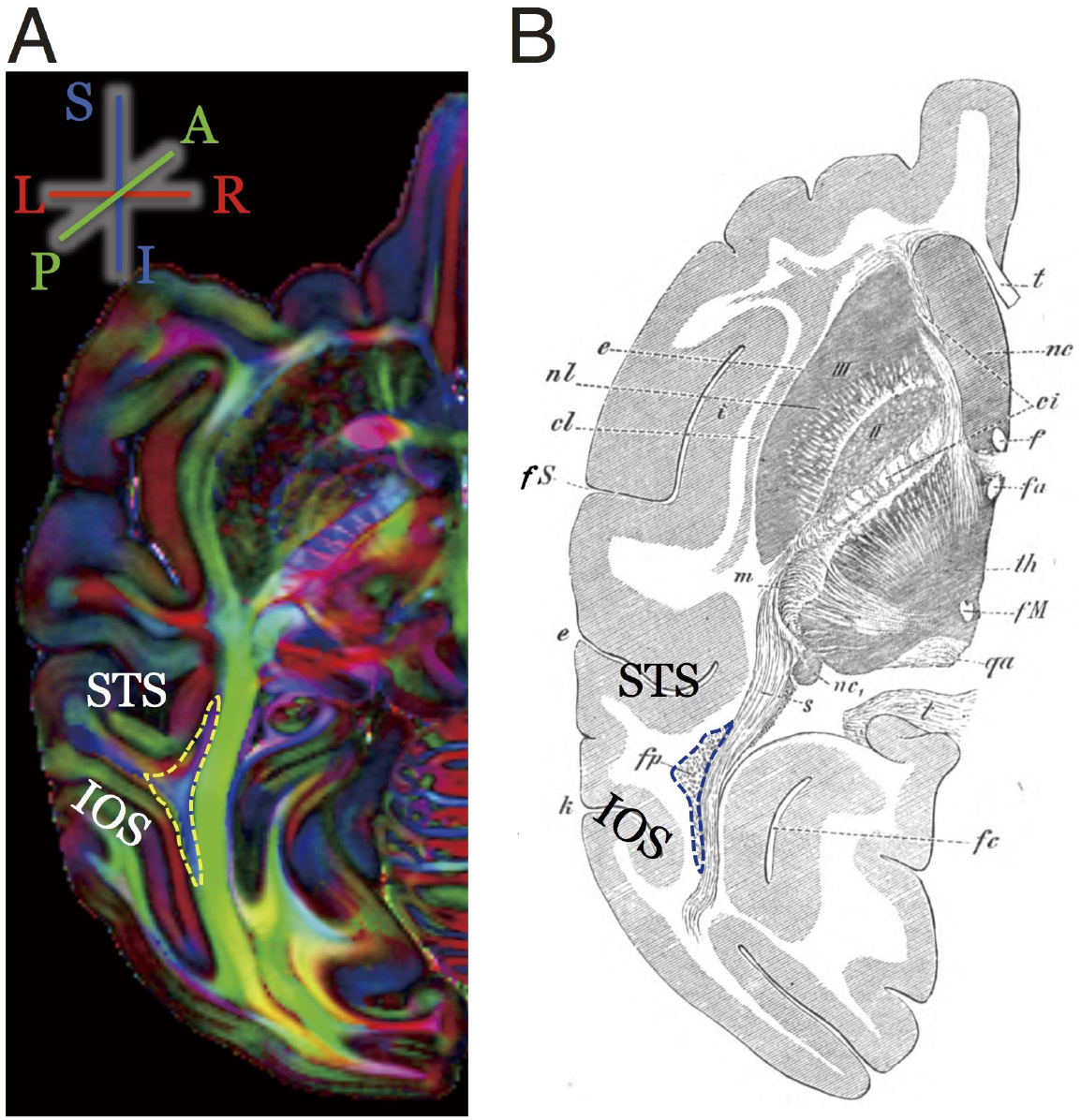
History of the macaque VOF. The comparison of the mVOF position in modern diffusion MRI data and Wernicke’s study (Wernicke 1881). **A.** The position of the mVOF in PDD map in the left hemisphere (subject M1). This slice is chosen to match the schematic diagram in Wernicke’s study (right, **B**). While it is impossible to completely match the slice between modern data and classical work, the position of the mVOF (left) and “perpendicular fasciculus” (*fp*, right panel) is qualitatively consistent; both are located between the STS (*e*, *Parallelfurche* in Wernicke) and IOS (*k, vordere Occipitalfurche* in Wernicke).

The dMRI derived mVOF is also consistent with tract degeneration studies conducted by Bailey and colleagues (Bailey et al. 1944). Specifically, they studied association pathways in *Macaca mulatta* by first injecting the neurotoxin in a target cortical site and then measuring the loss of electrophysiological signal following cortical damage. Using this method, Bailey and colleagues reported a pathway connecting the posterior end of the temporal cortex with both dorsal extrastriate cortex as well as the posterior portion of the inferior parietal lobule. Petr and colleagues (Petr et al. 1949) further explored these tracts with comparable methods. They identified that this vertical pathway extended from dorsal visual cortex and nearby aspects of the inferior parietal lobe to macaque TEO, but not area TE. This observation is consistent with results from the present study showing that the mVOF has ventral endpoints within TEO, but not TE (Figure 6). It is important to note that while the mVOF has not been widely examined in modern studies, these vertical connections were crucial for solidifying the division separating TEO from TE as these areas were complicated to differentiate cytoarchitectonically in Von Bonin and Bailey’s classic atlas (Iwai and Mishkin 1969; Petr et al. 1949).

Additionally, Bonin and colleagues (1942) provide observations that are consistent with the present work. For example, we find that the mVOF and ILF are compressed and intertwined within the occipital lobe of macaque, while the VOF and ILF are clearly separable with orthogonal orientations in human. Bonin and colleagues had very similar observations. In a summary of their strychnine measurements and Weigert stains in macaque, Bonin et al. write:

> Wernicke’s vertical bundle, running lateral of the ventricle, is well seen in the macaque wherein it was first recognized by Wernicke himself…there is much more room in the spacious white core of the occipital lobes of chimpanzee and man than in the sparse white matter of the macaque’s occipital lobe. Hence the association fibers which are compressed into a compact bundle in the macaque, may be more diffusely spread out over a larger space in the higher primates…The experimental results suggest moreover that there are fibers passing from the dorsal part of the parastriate area to the temporal lobe. These may run first in the vertical occipital bundle of Wernicke, to sweep over into the inferior horizontal fasciculus in a ventral level. It should not be too difficult to test this assumption. Pg. 179-183

The evidence from the present study provides support for their hypothesis nearly 75 years later.

Evidence from recent studies provide further anatomical evidence for the mVOF. For example, in recent studies, Schmahmann and Pandya explored major white matter tracts, including those in occipital cortex, in macaque brain using retrograde tracers (Schmahmann and Pandya 2006). Schmahmann and Pandya directly relate the ILF to vertical fascicles when they write:

> We have observed that fibers that are caudally situated in the vertical component of the ILF link the dorsal and ventral aspects of the occipital lobe and are equivalent to this vertical occipital system of Wernicke.

Their description of this fiber pathway is also consistent with what we observed in our dMRI data, showing the vertical occipital fiber connecting dorsal and ventral visual cortex which are less distinguishable from the ILF (Figure 7).

There is a collection of tracer studies of the macaque that are consistent with this study. Ungerleider and Desimone (1986) reported a connection between MT and ventral V4. Distler and colleagues (1993) injected anterograde and retrograde tracers in macaque area TEO, and described a bidirectional connections to dorsal areas, including V3d, V3A, V4d and MT. Webster and colleagues (Webster et al. 1994) explored the cortical connections in area TEO and TE, and reported that the TEO is connected with V3A whereas TE is predominantly connected to area LIPd, rather than V3A. More recently, Ungerleider and colleagues (Ungerleider et al. 2008) studied pathways terminating in macaque V4 using both anterograde and retrograde tracers. They described bidirectional connections between V3A, MT (dorsal) and ventral V4. These tracer studies suggest that the mVOF includes bidirectional information transmission between dorsal and ventral visual cortex. The streamline estimates here also show that the mVOF connects dorsal (V3A, V4d and MT) and ventral areas (V4v, TEO).

Taken together, the mVOF streamlines follow a path that is consistent with macaque studies using fiber dissections, tract degeneration, Weigert stains, and tracers.

### The VOF cortical endpoints with respect to visual field maps

The organization of the visual field map has been widely studied in both human and macaque system, and scientists have proposed organization principles of the visual field maps (Wandell et al. 2007). The VOF endpoints in human and macaque share some similarities with respect to the position of visual field maps. Notably, in both species, VOF endpoints are close to V3A dorsally and V4 ventrally. In both human fMRI and macaque electrophysiology literature, visual neuroscientists largely agree that relatively dorsal aspects of visual cortex process spatial information, whereas relatively ventral aspects of visual cortex process categorical information (Ungerleider and Mishkin 1982; Goodale and Milner 1992; Ungerleider and Haxby 1994). Recent studies raise the possibility of substantial interaction between dorsal and ventral visual areas (Grill-Spector et al. 1998, 2000; Konen and Kastner 2008). The increased prominence of the VOF in recent anatomy literature provides an anatomical infrastructure supporting the interaction between dorsal and ventral visual cortex (Yeatman et al. 2014; Takemura et al. 2016b).

In both human and macaque, V2 and V3 are separated into dorsal and ventral components, and each represent a quarter of the visual field. However, mid-level visual maps contain hemifield representations, such as V3A/B, IPS-0, hV4, VO, LO in humans and V3A in macaques. The macaque map organization anterior to V4v is controversial in the literature, but a recent study provides evidence for hemifield maps anterior to V4v and adjacent to TEO (Kolster et al. 2014). As endpoints of the VOF are near hemifield maps, it is likely that the VOF has an essential functional role for transmitting upper and lower visual field information between dorsal and ventral visual field maps (Takemura et al. 2016b).

While the VOF in human and macaque have some similarities, the human and macaque visual field maps have some differences in spatial arrangements (Wandell and Winawer 2011; Kolster et al. 2014; Vanduffel et al. 2014). For example, some differences between human and macaque reported in the literature are the volume of V3 (Brewer et al. 2002; Dougherty et al. 2003), the retinotopy in V4 (Brewer et al. 2005; Wade et al. 2008; Arcaro et al. 2009; Winawer et al. 2010; Goddard et al. 2011; Winawer and Witthoft 2015), and response selectivity in V3A (Tootell et al. 1997; Vanduffel et al. 2001). The homology of some humans maps, such as V3B (Smith et al. 1998; Press et al. 2001) and LO (Larsson and Heeger 2006; Amano et al. 2009; Silson et al., 2013), is still under debate (but see Kolster et al. 2014). The difference in the spatial arrangement of maps between human and macaque may be related to the change in position and size of the VOF. The position of all these maps, and their size, make it seem likely that the growth in the VOF and its change in position may have had influence on the cortical organization.

### The relative size and position of the VOF and ILF in human and macaque

Human and macaque VOF are similar in two critical ways. First, the endpoints of the VOF are near similar maps such as V3A in dorsal visual cortex and V4 in ventral visual cortex. Second, in both human and macaque, the VOF is located lateral to the Optic Radiation and oriented in a direction perpendicular to the OR. These similarities suggest that the mVOF is a homologous pathway to the human VOF. But there are also significant differences between these pathways that reflect some of the general differences between the macaque brain and the much larger human brain.

First, the human VOF is significantly elongated in the anterior-posterior dimension compared to the macaque VOF, which is largely confined between the IOS and STS. The human VOF extends nearly to the posterior Arcuate Fasciculus (pAF; or Vertical (posterior) Segment of the Arcuate; Catani et al. 2005; Weiner et al. 2016b). While classical and recent studies describe the pAF and VOF as separate tracts (Curran 1909; Catani et al. 2005; Yeatman et al. 2014; Weiner et al. 2016b), the distinction between the VOF and pAF is relatively unclear in the literature. As discussed in a recent letter (Weiner et al. 2016b), there is uncertainty regarding the homology between mVOF relative to the human VOF and pAF for a number of reasons. For example, the human brain is significantly expanded compared to the macaque brain and there is not a one-to-one mapping of anatomical structures between species. Consequently, mapping Wernicke’s original definition of the perpendicular fasciculus in a schematic image to the human brain is non-trivial.

In fact, some groups suggest that the pAF is homologous to mVOF (also termed Wernicke’s perpendicular fasciculus; Homola et al. 2012; Bartsch et al. 2013; see also Martino and Garcia-Porrero 2013 for a debate). Our comparison with visual maps suggests that the mVOF is homologous to human VOF, because it has endpoints near similar maps such as V3A and ventral V4 (Figure 6). While there are significant similarities between human and macaque VOF, the relation of human pAF with mVOF is an open question. One possibility is that the pAF and the VOF in humans originally derived from the same vertical fiber system in a common ancestor between humans and macaques. Throughout the evolution and expansion of frontal cortex and white matter pathways, the vertical fiber system of the human brain may have expanded in the anterior-posterior direction whereby the most anterior fascicles became the pAF in human.

A second difference is that the macaque mVOF and ILF streamlines intermingle, but in human the estimated VOF streamlines are largely separate and lateral to the estimated ILF streamlines. This difference suggests a hypothesis regarding the evolution of the VOF that is related to the substantial cortical expansion in human (Figure 11). The VOF may be the part of the ILF system in a common ancestor (right panel, Figure 10). With the expansion of cerebral cortex and the increase in the number of mid-level visual areas, the VOF may have grown to the point where it became independent of the ILF system, shifting laterally and separating from the ILF. This hypothesis is speculative because it is difficult to exclude the possibility that the estimated human ILF streamlines separate from the VOF due to the limited ability to resolve crossing fibers in *in vivo* human data or waypoint ROI selection procedures. We include this speculation because it provides a hypothesis for future comparative studies, comparing high-resolution human and macaque anatomy data, that assess the evolution of the visual association areas and vertical occipital fiber system in more fine detail.

**Figure 11.**
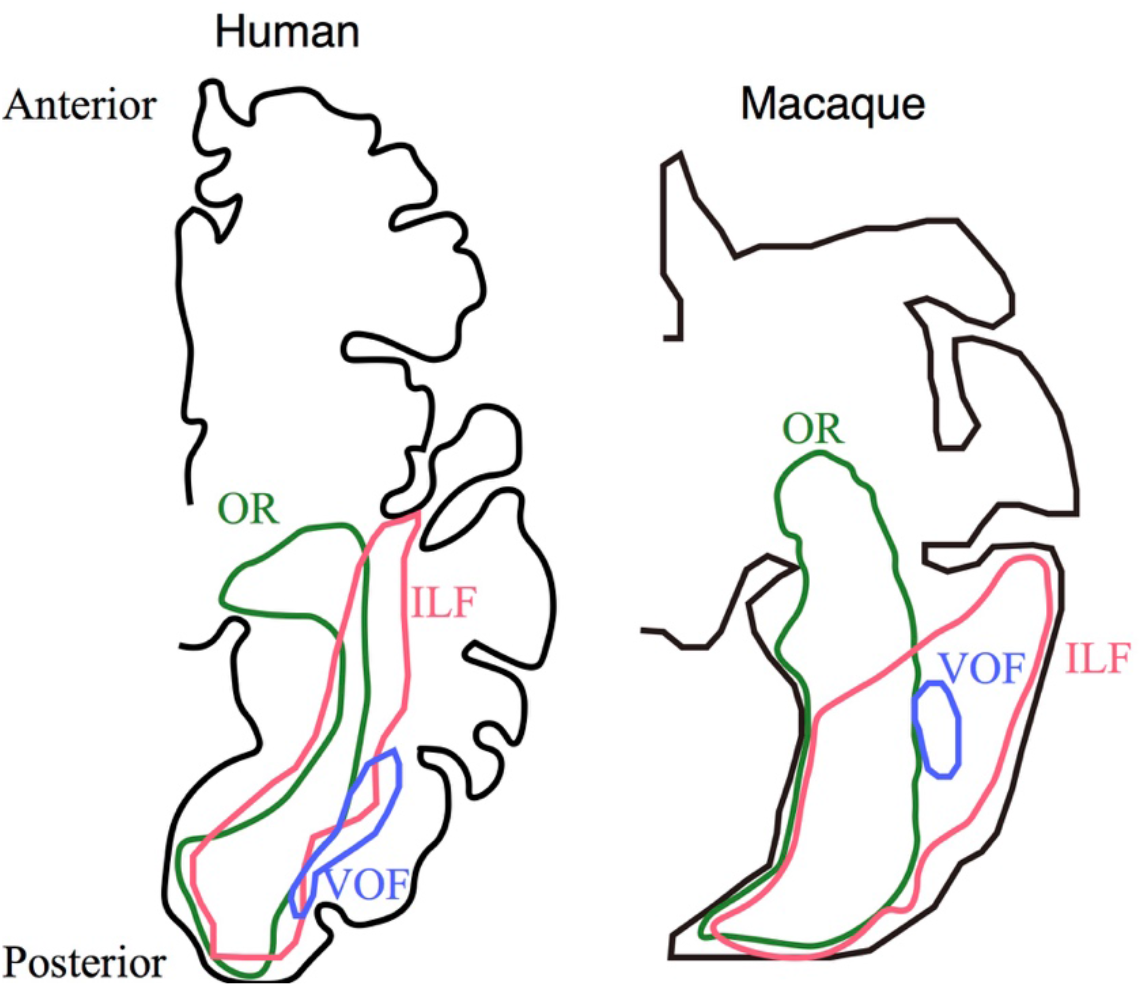
Schematic diagram of estimated human-macaque occipital fiber system. This diagram schematically describes the position of three major white matter pathways in the right hemisphere in humans (left panel) and macaque (right panel); the OR (green), the ILF (red) and the VOF (blue). As compared with macaques, human VOF is moved to the lateral side of the white matter and becomes relatively distinguishable from the ILF.

### Combining post-mortem anatomical data with *in vivo* diffusion MRI

Tractography creates a model of the white matter tracts, and the model is constrained to predict the diffusion MRI data. Current models and data are generally at the millimeter scale and decreasing. But even the highest quality current measurements do not provide accurate estimates on cortico-cortical connectivity at the spatial scale of fine connections (Thomas et al. 2014; Reveley et al. 2015). It is worth remembering, however, that much information can be gleaned from millimeter scale measurements in living subjects, and we should take advantage of what we can learn from this scale.

The data in this paper include *in vivo* diffusion data and post-mortem diffusion data, and we contextualize the diffusion measurements with post-mortem tracer studies. The in vivo diffusion data are relatively coarse, but they have great value for studies of development, plasticity, disease and individual differences. This type of data is being used in human studies to provide insight of white matter plasticity during developmental stages in relation to behavioral learning (Yeatman et al. 2012a; Gomez et al. 2015) or white matter consequences of retinal or cortical damages (Levin et al. 2010; Bock et al. 2013; Ogawa et al. 2014; Ajina et al. 2015). Recent studies use these methods to measure tissue changes following learning and disease in rodents (Blumenfeld-Katzir et al. 2011; Hofstetter et al. 2013; Sampaio-Baptista et al. 2013), marmosets (Warner et al. 2015) and macaques (Li et al. 2011).

There are several advantages of *in vivo* dMRI compared to other methods. First, *in vivo* methods enable longitudinal measurements. Second, dMRI provides a large field of view compared to post-mortem tracer studies. Third, dMRI and tractography have built up an infrastructure of reproducible computational methods and atlases that can be shared (Hayashi et al., 2015; Calabrese et al. 2015). This supports the methods needed to grow a shared database that enables comparisons between individual subjects and population norms (Wandell et al. 2015).

The strengths and weaknesses of anatomical methods are complementary to the dMRI strengths. Post-mortem tracer studies are very well suited for examining hypotheses about the fine spatial detail and directionality of specific connections (Cheng et al., 1997; Ungerleider et al. 2008; Banno et al., 2011; Grimaldi et al. 2016), or the relationship between specific connections and molecular targets (Ichinohe et al. 2010). For example, if one identifies a target of interest using dMRI, the best way to discover the molecular and cellular basis is to make anatomical and histological measurements in locations identified as good targets from the dMRI data (Sagi et al. 2012; Sampaio-Baptista et al. 2013). In contrast, it is very difficult to study the variability between individuals using tracer methods because of the inability to precisely control the placement of injections and uptake of tracers. Anatomical tracer studies typically slice the brain and recover only a subset of the pathways reconstructed from challenging material (Kennedy et al. 2013). As a result, these studies have a limited field of view and critical measurement limitations to the result from the need to dissect the brain to follow tracts. Finally, tracer studies have not yet developed a digital technology that enables one to compute atlases and integrate data. The most widely used summaries of anatomical data are simple tables that collect the known collection, and these tables do not capture the pathways themselves (CoCoMac database; Bakker et al. 2012). The methods for integrating anatomical data with diffusion involves comparing drawings from tracer studies with digital atlases in a reference coordinate frame and reproducible computations from dMRI studies.

Once we recognize that tractography is model building, it becomes clear that there is great potential value in combining post-mortem and *in vivo* dMRI data. Both measure a large field of view and can be represented in a computational format. The post-mortem studies obtain data and build models at higher resolution and better sensitivity. These models can serve to disambiguate the lower quality *in vivo* measurements. Moreover, because the data are obtained without slicing the brain the post-mortem data are easily interpreted in terms of the natural three-dimensional pathway structures.

Finally, when high quality post-mortem measurements provide a consistent and high quality model (Axer et al. 2011; Miller et al. 2011; Dell’Acqua et al. 2013; Leuze et al. 2014; Aggarwal et al. 2015; Caspers et al., 2015; Seehaus et al. 2015), the same model can guide the search for tracts using the lower resolution *in vivo* dMRI data. In this approach, the post-mortem data are used to create high quality tract models. The *in vivo* data are used to find the most likely position of known tracts (Sherbondy et al. 2008b; Kammen et al., 2016). To some extent, this has been the approach used in much of human tractography. The major white matter tracts that are studied by many groups are known to exist, and tractography is used to identify their positions and paths (Catani et al. 2002; Wakana et al. 2004; Catani and Thiebaut de Schotten 2008; Yendiki et al. 2011; Yeatman et al. 2012b).

In summary, one way to proceed is to develop high-resolution models of the white matter pathways using post-mortem diffusion data, and to use these models to guide model estimates from lower resolution *in vivo* data. We can then use anatomical tracer methods to analyze specific pathways and cellular molecular questions.

### Conclusions

The comparative analysis reported here supports the long-held view that certain aspects of human and macaque visual cortex are very similar. Specifically, the majority of occipital white matter tracts (OR, Forceps Major, ILF) appear to form similar connections in the two species. Significant differences, on the other hand, were found in the white matter tract connecting occipital and frontal cortex. Most significant is the absence in macaque of a tract homologous to human IFOF. The much larger volume (15x) of the human brain seems to have brought forth a need to create a long pathway that speeds the projection of signals from posterior to anterior portions of the brain. Such a new pathway together with the well-documented expansion of the frontal cortex in humans appears to have brought for new circuitry underlying visual cognitive behavior. Tasks that rely on this communication pathway in human may not have a corresponding substrate in macaque.

A second notable difference is the position of the mVOF and the human VOF with respect to other white matter tracts. In both species, the VOF connects dorsal and ventral posterior brain, and are likely to have endpoints near similar maps (V3A and ventral V4). But, in human, the VOF is largely segregated from the ILF while in macaque, the VOF is embedded within the ILF. This difference will certainly have implications for interpreting the consequences of white matter lesions, such as one might observe in stroke patients or other clinical lesions. It is not clear that the position matters for typical signaling and function.

The gross anatomical positions of stimulus-selective cortical regions, such as regions selective for faces, differ between human and macaque (Tsao et al. 2008; Bell et al. 2009; Pinsk et al. 2009; Kornblith et al. 2013; Weiner and Grill-Spector 2015; Lafer-Sousa et al. 2016). The dMRI measurements reveal similarities and differences in the organization of the major occipital white matter tracts in human and macaque, and the projections of these tracts may explain the different locations of these cortical specializations. It is possible that in some cases, the different position is incidental, and the fundamental white matter circuitry is homologous. As we improve our understanding about the white matter tracts in both species, we hope to clarify the degree to which certain cortical specializations reflect a common functional circuitry in which responses in macaque serve as an accurate model for human vision.

## Acknowledgements

We thank Ariel Rokem and Jason D. Yeatman for supporting in the collection of human diffusion MRI data, Stelios Smirnakis for supporting in the collection of macaque diffusion MRI data, Kaoru Amano, Atsushi Wada and Noboru Nushi for providing the computer environment, Cesar F. Caiafa for providing analysis tools, and Lee Michael Perry for technical assistance. This study is funded in part by JSPS Postdoctoral Fellowship for Research Abroad and the Grant-in-Aid for JSPS Research Fellow (to H.T.), Indiana University College of Arts and Sciences startup funds (to F.P.), Indiana Clinical and Translational Sciences Institute CTSI (GLUE Grant; supported by NIH grants ULTTR001108, ULTTR001106, ULTTR001107; to F.P.), Human Frontier Science Program Long-Term Fellowship (LT000418/2013-L, to J.S.), a Fondation pour la Recherche Médicale Post-doctoral fellowship (to J.S.), a Women&Science Post-doctoral Fellowship (to J.S.), a Bettencourt-Schueller Foundation Young Researcher Award (to J.S.), the intramural Research Program of the National Institutes of Health (to F.Q.Y. and D.A.L.), a Pew Scholar Award in the Biomedical Sciences (to W.A.F.), The Esther A. & Joseph Klingenstein Fund (to W.A.F.), a McKnight Scholars Award (to W.A.F.), the New York Stem Cell Foundation (NYSCF-R-NI23, to W.A.F.), the National Eye Institute (R01 EY021594-01A1, to W.A.F.) and NSF BCS-1228397 (to B.A.W.). H.T. is a Superlative Postdoctoral Fellow of Japan Society for the Promotion of Science. S.M.L. is a Howard Hughes Medical Institute International Student Research fellow. W.A.F. is a New York Stem Cell Foundation-Robertson Investigator. Data were provided in part by the Human Connectome Project, WU-Minn Consortium (Van Essen, D. and Ugurbil, K., 1U54MH091657).

## References

Aggarwal M, Nauen DW, Troncoso JC, Mori S. 2015. Probing region-specific microstructure of human cortical areas using high angular and spatial resolution diffusion MRI. Neuroimage. 105:198–207.

Ajina S, Pestilli F, Rokem A, Kennard C, Bridge H. 2015. Human blindsight is mediated by an intact geniculo-extrastriate pathway. eLife. 4:e08935.

Amano K, Wandell BA, Dumoulin SO. 2009. Visual field maps, population receptive field sizes, and visual field coverage in the human MT+ complex. J Neurophysiol. 102:2704–2718.

Arcaro MJ, McMains SA, Singer BD, Kastner S. 2009. Retinotopic organization of human ventral visual cortex. J Neurosci. 29:10638–10652.

Axer M, Amunts K, Grässel D, Palm C, Dammers J, Axer H, Pietrzyk U, Zilles K. 2011. A novel approach to the human connectome: ultra-high resolution mapping of fiber tracts in the brain. Neuroimage. 54:1091–1101.

Bailey P, Von Bonin G, Davis EW, Garol HW, Mcculloch WS. 1944. Further observatioms on associational pathways in the brain of macaca mulatta. J Neuropathol Exp Neurol. 3:413.

Bakker R, Wachtler T, Diesmann M. 2012. CoCoMac 2.0 and the future of tract-tracing databases. Front Neuroinform. 6:30.

Banno T, Ichinohe N, Rockland KS, Komatsu H. 2011. Reciprocal connectivity of identified color-processing modules in the monkey inferior temporal cortex. Cereb Cortex. 21:1295–1310.

Bartsch AJ, Jbabdi S, Geletneky K. 2013. The Temporo-parietal Fiber Intersection Area (TPFIA) and Wernicke’s Perpendicular Fasciculus (WpF). Neurosurgery. 73:E381–2.

Bell AH, Hadj-Bouziane F, Frihauf JB, Tootell RBH, Ungerleider LG. 2009. Object representations in the temporal cortex of monkeys and humans as revealed by functional magnetic resonance imaging. J Neurophysiol. 101:688–700.

Berman JI, Lanza MR, Blaskey L, Edgar JC, Roberts TPL. 2013. High angular resolution diffusion imaging probabilistic tractography of the auditory radiation. AJNR Am J Neuroradiol. 34:1573–1578.

Blumenfeld-Katzir T, Pasternak O, Dagan M, Assaf Y. 2011. Diffusion MRI of structural brain plasticity induced by a learning and memory task. PLoS One. 6:e20678.

Bock AS, Saenz M, Tungaraza R, Boynton GM, Bridge H, Fine I. 2013 Visual callosal topography in the absence of retinal input. Neuroimage. 81:325–334.

Brewer AA, Liu J, Wade AR, Wandell BA. 2005. Visual field maps and stimulus selectivity in human ventral occipital cortex. Nat Neurosci. 8:1102–1109.

Brewer AA, Press WA, Logothetis NK, Wandell BA. 2002. Visual areas in macaque cortex measured using functional magnetic resonance imaging. J Neurosci. 22:10416–10426.

Bullock TH, Bennett MV, Johnston D, Josephson R, Marder E, Fields RD. 2005. Neuroscience. The neuron doctrine, redux. Science. 310:791–793.

Caiafa CF, Pestilli F. 2015. Sparse multiway decomposition for analysis and modeling of diffusion imaging and tractography. ArXiv. 1505.0710.

Calabrese E, Badea A, Coe CL, Lubach GR, Shi Y, Styner MA, Johnson GA. 2015. A diffusion tensor MRI atlas of the postmortem rhesus macaque brain. Neuroimage. 117:408–416.

Caspers S, Axer M, Caspers J, Jockwitz C, Jütten K, Reckfort J, Grässel D, Amunts K, Zilles K. 2015. Target sites for transcallosal fibers in human visual cortex - A combined diffusion and polarized light imaging study. Cortex. 72:40–53.

Catani M, Thiebaut de Schotten M. 2012. Atlas of human brain connections. Oxford University Press.

Catani M, Ffytche DH. 2005. The rises and falls of disconnection syndromes. Brain. 128:2224–2239.

Catani M, Howard RJ, Pajevic S, Jones DK. 2002. Virtual in vivo interactive dissection of white matter fasciculi in the human brain. Neuroimage. 17:77–94.

Catani M, Jones DK, Ffytche DH. 2005. Perisylvian language networks of the human brain. Ann Neurol. 57:8–16.

Catani M, Thiebaut de Schotten M. 2008. A diffusion tensor imaging tractography atlas for virtual in vivo dissections. Cortex. 44:1105–1132.

Cheng K, Saleem KS, Tanaka K. 1997. Organization of corticostriatal and corticoamygdalar projections arising from the anterior inferotemporal area TE of the macaque monkey: a Phaseolus vulgaris Leucoagglutinin Study. J Neurosci. 17:7902–7295.

Corthout EC. 2014. The eye and brain in macaque and man: linear, areal and volumetric dimensions. Leuven, Belgium: ACCO.

Craddock RC, Jbabdi S, Yan CG, Vogelstein JT, Castellanos FX, Di Martino A, Kelly C, Heberlein K, Colcombe S, Milham MP. 2013. Imaging human connectomes at the macroscale. Nat Methods. 10:524–539.

Curran EJ. 1909. A new association fiber tract in the cerebrum with remarks on the fiber tract dissection method of studying the brain. J Comp Neurol Psychol. 19:645–656.

Déjerine J. 1895. Anatomie des centres nerveux. Paris: Rueff et Cie.

Dell’Acqua F, Bodi I, Slater D, Catani M, Modo M. 2013. MR diffusion histology and micro-tractography reveal mesoscale features of the human cerebellum. Cerebellum. 12:923–931.

De Valois RL, Jacobs GH. 1968. Primate color vision. Science. 162:533–540.

De Valois RL, Morgan HC, Polson MC, Mead WR, Hull EM. 1974. Psychophysical studies of monkey vision—I. Macaque luminosity and color vision tests. Vision Res. 14:53–67.

Distler C, Boussaoud D, Desimone R, Ungerleider LG. 1993. Cortical connections of inferior temporal area TEO in macaque monkeys. J Comp Neurol. 334:125–150.

Dougherty RF, Koch VM, Brewer AA, Fischer B, Modersitzki J, Wandell BA. 2003. Visual field representations and locations of visual areas V1/2/3 in human visual cortex. J Vis. 3:586–598.

Duan Y, Norcia AM, Yeatman JD, Mezer A. 2015. The structural properties of major white matter tracts in strabismic amblyopia. Invest Ophthalmol Vis Sci. 56:5152–5160.

Duffau H, Herbet G, Moritz-Gasser S. 2013. Toward a pluri-component, multimodal, and dynamic organization of the ventral semantic stream in humans: lessons from stimulation mapping in awake patients. Front Syst Neurosci. 7:44.

Dumoulin SO, Wandell BA. 2008. Population receptive field estimates in human visual cortex. Neuroimage. 39:647–660.

Fields RD. 2008a. White matter matters. Sci Am. 298:42–49.

Fields RD. 2008b. White matter in learning, cognition and psychiatric disorders. Trends Neurosci. 31:361–370.

Fields RD. 2015. A new mechanism of nervous system plasticity: activity-dependent myelination. Nat Rev Neurosci. 16:756–767.

Fischl B. 2012. FreeSurfer. Neuroimage. 62:774–781.

Fize D, Vanduffel W, Nelissen K, Denys K, Chef d’Hotel C, Faugeras O, Orban GA. 2003. The retinotopic organization of primate dorsal V4 and surrounding areas: A functional magnetic resonance imaging study in awake monkeys. J Neurosci. 23:7395–7406.

Forkel SJ, Thiebaut de Schotten M, Kawadler JM, Dell’Acqua F, Danek A, Catani M. 2014. The anatomy of fronto-occipital connections from early blunt dissections to contemporary tractography. Cortex. 56:73–84.

Friston KJ, Ashburner J. 2004. Generative and recognition models for neuroanatomy. Neuroimage. 23:21–24.

Gellman RS, Carl JR, Miles FA. 1990. Short latency ocular-following responses in man. Vis Neurosci. 5:107–122.

Goda N, Tachibana A, Okazawa G, Komatsu H. 2014. Representation of the material properties of objects in the visual cortex of nonhuman primates. J Neurosci. 34:2660–2673.

Goddard E, Mannion DJ, McDonald JS, Solomon SG, Clifford CW. 2011. Color responsiveness argues against a dorsal component of human V4. J Vis. 11(4):1–21.

Gomez J, Pestilli F, Witthoft N, Golarai G, Liberman A, Poltoratski S, Yoon J, Grill-Spector K. 2015. Functionally defined white matter reveals segregated pathways in human ventral temporal cortex associated with category-specific processing. Neuron. 85:216–227.

Goodale MA, Milner AD. 1992. Separate visual pathways for perception and action. Trends Neurosci. 15:20–25.

Greenberg AS, Verstynen T, Chiu YC, Yantis S, Schneider W, Behrmann M. 2012. Visuotopic cortical connectivity underlying attention revealed with white-matter tractography. J Neurosci. 32:2773–2782.

Grill-Spector K, Kushnir T, Edelman S, Itzchak Y, Malach R. 1998. Cue-invariant activation in object-related areas of the human occipital lobe. Neuron. 21:191–202.

Grill-Spector K, Kushnir T, Hendler T, Malach R. 2000. The dynamics of object-selective activation correlate with recognition performance in humans. Nat Neurosci. 3:837–843.

Grimaldi P, Saleem KS, Tsao D. 2016. Anatomical connections of the functionally defined “face patches” in the macaque monkey. Neuron. 90:1325–1342.

Hayashi T, Zhang G, Urayama S, Ose T, Watabe H, Onoe K, Tanki N, Murata Y, Higo N, Onoe H. 2015. High-resolution diffusion and structural MRI brain atlas of rhesus macaques. In: Organization for the Human Brain Mapping Annual Meeting, Honolulu. HI.

Hinds OP, Rajendran N, Polimeni JR, Augustinack JC, Wiggins G, Wald LL, Diana Rosas H, Potthast A, Schwartz EL, Fischl B. 2008. Accurate prediction of V1 location from cortical folds in a surface coordinate system. Neuroimage. 39:1585–1599.

Hofstetter S, Tavor I, Tzur Moryosef S, Assaf Y. 2013. Short-term learning induces white matter plasticity in the fornix. J Neurosci. 33:12844–12850.

Homola GA, Jbabdi S, Beckmann CF, Bartsch AJ. 2012. A brain network processing the age of faces. PLoS One. 7:e49451.

Honey CJ, Sporns O. 2008. Dynamical consequences of lesions in cortical networks. Hum Brain Mapp. 29:802–809.

Horwitz GD. 2015. What studies of macaque monkeys have told us about human color vision. Neuroscience. 296:110–115.

Ichinohe N, Matsushita A, Ohta K, Rockland KS. 2010. Pathway-specific utilization of synaptic zinc in the macaque ventral visual cortical areas. Cereb Cortex. 20:2818–2831.

Iturria-Medina Y, Perez Fernandez A, Morris DM, Canales-Rodriguez EJ, Haroon HA, Garcia Penton L, Augath M, Galan Garcia L, Logothetis N, Parker GJ, Melie-Garcia L. 2011. Brain hemispheric structural efficiency and interconnectivity rightward asymmetry in human and nonhuman primates. Cereb Cortex. 21:56–67.

Iwai E, Mishkin M. 1969/9. Further evidence on the locus of the visual area in the temporal lobe of the monkey. Exp Neurol. 25:585–594.

Jbabdi S, Lehman JF, Haber SN, Behrens TE. 2013. Human and monkey ventral prefrontal fibers use the same organizational principles to reach their targets: tracing versus tractography. Journal of Neuroscience. 33:3190–3201.

Kammen A, Law M, Tjan BS, Toga AW, Shi Y. 2016. Automated retinofugal visual pathway reconstruction with multi-shell HARDI and FOD-based analysis. Neuroimage. 125:767–779.

Kennedy H, Knoblauch K, Toroczkai Z. 2013. Why data coherence and quality is critical for understanding interareal cortical networks. Neuroimage. 80:37–45.

Kim M, Ronen I, Ugurbil K, Kim D-S. 2006. Spatial resolution dependence of DTI tractography in human occipito-callosal region. Neuroimage. 32:1243–1249.

Kolster H, Janssens T, Orban GA, Vanduffel W. 2014. The retinotopic organization of macaque occipitotemporal cortex anterior to V4 and caudoventral to the middle temporal (MT) cluster. J Neurosci. 34:10168–10191.

Konen CS, Kastner S. 2008. Two hierarchically organized neural systems for object information in human visual cortex. Nat Neurosci. 11:224–231.

Kornblith S, Cheng X, Ohayon S, Tsao DY. 2013. A network for scene processing in the macaque temporal lobe. Neuron. 79:766–781.

Kriegeskorte N, Mur M, Ruff DA, Kiani R, Bodurka J, Esteky H, Tanaka K, Bandettini PA. 2008. Matching categorical object representations in inferior temporal cortex of man and monkey. Neuron. 60:1126–1141.

Lafer-Sousa R, Conway BR, Kanwisher NG. 2016. Color-biased regions of the ventral visual pathway lie between face-and place-selective regions in humans, as in macaques. J Neurosci. 36:1682–1697.

Larsson J, Heeger DJ. 2006. Two retinotopic visual areas in human lateral occipital cortex. J Neurosci. 26:13128–13142.

Lebel C, Benner T, Beaulieu C. 2012. Six is enough? Comparison of diffusion parameters measured using six or more diffusion-encoding gradient directions with deterministic tractography. Magn Reson Med. 68:474–483.

Lee JH, Garwood M, Menon R, Adriany G, Andersen P, Truwit CL, Ugurbil K. 1995. High contrast and fast three-dimensional magnetic resonance imaging at high fields. Magn Reson Med. 34:308–312.

Leong JK, Pestilli F, Wu CC, Samanez-Larkin GR, Knutson B. 2016. White-matter tract connecting anterior insula to nucleus accumbens correlates with reduced preference for positively skewed gambles. Neuron. 89:63–69.

Leuze CW, Anwander A, Bazin PL, Dhital B, Stuber C, Reimann K, Geyer S, Turner R. 2014. Layer-specific intracortical connectivity revealed with diffusion MRI. Cereb Cortex. 24:328–339.

Levin N, Dumoulin SO, Winawer J, Dougherty RF, Wandell BA. 2010. Cortical maps and white matter tracts following long period of visual deprivation and retinal image restoration. Neuron. 65:21–31.

Li C, Zhang X, Komery A, Li Y, Novembre FJ, Herndon JG. 2011. Longitudinal diffusion tensor imaging and perfusion MRI investigation in a macaque model of neuro-AIDS: a preliminary study. Neuroimage. 58:286–292.

Li L, Hu X, Preuss TM, Glasser MF, Damen FW, Qiu Y, Rilling J. 2013. Mapping putative hubs in human, chimpanzee and rhesus macaque connectomes via diffusion tractography. Neuroimage. 80:462–474.

Lindbloom-Brown Z, Tait LJ, Horwitz GD. 2014. Spectral sensitivity differences between rhesus monkeys and humans: implications for neurophysiology. J Neurophysiol. 112:3164–3172.

Lyon DC, Connolly JD. 2012. The case for primate V3. Proc Biol Sci. 279:625–633.

Mantini D, Corbetta M, Romani GL, Orban GA, Vanduffel W. 2012. Data-driven analysis of analogous brain networks in monkeys and humans during natural vision. Neuroimage. 63:1107–1118.

Mars RB, Foxley S, Verhagen L, Jbabdi S, Sallet J, Noonan MP, Neubert F-X, Andersson JL, Croxson PL, Dunbar RIM, Khrapitchev AA, Sibson NR, Miller KL, Rushworth MFS. 2015. The extreme capsule fiber complex in humans and macaque monkeys: a comparative diffusion MRI tractography study. Brain Struct Funct. [Epub Ahead of Print]

Martino J, Brogna C, Robles SG, Vergani F, Duffau H. 2010a. Anatomic dissection of the inferior fronto-occipital fasciculus revisited in the lights of brain stimulation data. Cortex. 46:691–699.

Martino J, Vergani F, Robles SG, Duffau H. 2010b. New insights into the anatomic dissection of the temporal stem with special emphasis on the inferior fronto-­-occipital fasciculus: implications in surgical approach. Neurosurgery. 66:ons4–ons12.

Martino J, De Witt Hamer PC, Vergani F, Brogna C, de Lucas EM, Vazquez-Barquero A, Garcia-Porrero JA, Duffau H. 2011. Cortex-sparing fiber dissection: an improved method for the study of white matter anatomy in the human brain. J Anat. 219:531–541.

Martino J, Garcia-Porrero JA. 2013. In Reply: Wernicke’s Perpendicular Fasciculus and Vertical Portion of the Superior Longitudinal Fasciculus. Neurosurgery. 73:E382–3.

Miller KL, Stagg CJ, Douaud G, Jbabdi S, Smith SM, Behrens TEJ, Jenkinson M, Chance SA, Esiri MM, Voets NL, Jenkinson N, Aziz TZ, Turner MR, Johansen-Berg H, McNab JA. 2011. Diffusion imaging of whole, post-mortem human brains on a clinical MRI scanner. Neuroimage. 57:167–181.

Miura K, Matsuura K, Taki M, Tabata H, Inaba N, Kawano K, Miles FA. 2006. The visual motion detectors underlying ocular following responses in monkeys. Vision Res. 46:869–878.

Mori S, Zhang J. 2006. Principles of diffusion tensor imaging and its applications to basic neuroscience research. Neuron. 51:527–539.

Newsome WT, Britten KH, Movshon JA. 1989. Neuronal correlates of a perceptual decision. Nature. 341:52–54.

Ogawa S, Takemura H, Horiguchi H, Terao M, Haji T, Pestilli F, Yeatman JD, Tsuneoka H, Wandell BA, Masuda Y. 2014. White matter consequences of retinal receptor and ganglion cell damage. Invest Ophthalmol Vis Sci. 55:6976–6986.

Okazawa G, Goda N, Komatsu H. 2012. Selective responses to specular surfaces in the macaque visual cortex revealed by fMRI. Neuroimage. 63:1321–1333.

Orban GA, Van Essen D, Vanduffel W. 2004. Comparative mapping of higher visual areas in monkeys and humans. Trends Cogn Sci. 8:315–324.

Pajevic S, Pierpaoli C. 1999. Color schemes to represent the orientation of anisotropic tissues from diffusion tensor data: application to white matter fiber tract mapping in the human brain. Magn Reson Med. 42:526–540.

Passingham R. 2009. How good is the macaque monkey model of the human brain? Curr Opin Neurobiol. 19:6–11.

Pestilli F, Yeatman JD, Rokem A, Kay KN, Wandell BA. 2014. Evaluation and statistical inference for human connectomes. Nat Methods. 11:1058–1063.

Petr R, Holden LB, Jirout J. 1949. The efferent intercortical connectionss of the surperficial corten of the temporal lobe (macaca mulatta)*. J Neuropathol Exp Neurol. 8:100.

Pierpaoli C, Walker L, Irfanoglu MO, Barnett A, Basser PJ, Chang L-C, Koay C, Pajevic S, Rohde G, Sarlls J, Wu M. 2010. TORTOISE: an integrated software package for processing of diffusion MRI data. In: ISMRM 18th annual meeting. Stockholm, Sweden.

Pinsk MA, Arcaro M, Weiner KS, Kalkus JF, Inati SJ, Gross CG, Kastner S. 2009. Neural representations of faces and body parts in macaque and human cortex: a comparative FMRI study. J Neurophysiol. 101:2581–2600.

Polosecki P, Moeller S, Schweers N, Romanski LM, Tsao DY, Freiwald WA. 2013. Faces in motion: selectivity of macaque and human face processing areas for dynamic stimuli. J Neurosci. 33:11768–11773.

Press WA, Brewer AA, Dougherty RF, Wade AR, Wandell BA. 2001. Visual areas and spatial summation in human visual cortex. Vision Res. 41:1321–1332.

Rajalingham R, Schmidt K, DiCarlo JJ. 2015. Comparison of object recognition behavior in human and monkey. J Neurosci. 35:12127–12136.

Reese TG, Heid O, Weisskoff RM, Wedeen VJ. 2003. Reduction of eddy-current-induced distortion in diffusion MRI using a twice-refocused spin echo. Magn Reson Med. 49:177–182.

Reveley C, Seth AK, Pierpaoli C, Silva AC, Yu D, Saunders RC, Leopold DA, Ye FQ. 2015. Superficial white matter fiber systems impede detection of long-range cortical connections in diffusion MR tractography. Proc Natl Acad Sci U S A. 112:E2820–E2828.

Rilling JK. 2006. Human and nonhuman primate brains: are they allometrically scaled versions of the same design? Evol Anthropol. 15:65–77.

Rilling JK, Glasser MF, Preuss TM, Ma X, Zhao T, Hu X, Behrens TEJ. 2008. The evolution of the arcuate fasciculus revealed with comparative DTI. Nat Neurosci. 11:426–428.

Roebroeck A, Galuske R, Formisano E, Chiry O, Bratzke H, Ronen I, Kim D-S, Goebel R. 2008. High-resolution diffusion tensor imaging and tractography of the human optic chiasm at 9.4 T. Neuroimage. 39:157–168.

Rohde GK, Barnett AS, Basser PJ, Marenco S, Pierpaoli C. 2004. Comprehensive approach for correction of motion and distortion in diffusion-weighted MRI. Magn Reson Med. 51:103–114.

Rokem A, Yeatman JD, Pestilli F, Kay KN, Mezer A, van der Walt S, Wandell BA. 2015. Evaluating the accuracy of diffusion MRI models in white matter. PLoS One. 10:e0123272.

Russ BE, Leopold DA. 2015. Functional MRI mapping of dynamic visual features during natural viewing in the macaque. Neuroimage. 109:84–94.

Sagi Y, Tavor I, Hofstetter S, Tzur-Moryosef S, Blumenfeld-Katzir T, Assaf Y. 2012. Learning in the fast lane: new insights into neuroplasticity. Neuron. 73:1195–1203.

Saleem KS, Logothetis NK. 2012. A combined MRI and histology atlas of the rhesus monkey brain in stereotaxic coordinates. San Diego: Academic Press.

Sampaio-Baptista C, Khrapitchev AA, Foxley S, Schlagheck T, Scholz J, Jbabdi S, DeLuca GC, Miller KL, Taylor A, Thomas N, Kleim J, Sibson NR, Bannerman D, Johansen-Berg H. 2013. Motor skill learning induces changes in white matter microstructure and myelination. J Neurosci. 33:19499–19503.

Sarubbo S, De Benedictis A, Maldonado IL, Basso G, Duffau H. 2013. Frontal terminations for the inferior fronto-occipital fascicle: anatomical dissection, DTI study and functional considerations on a multi-component bundle. Brain Struct Funct. 218:21–37.

Sasaki Y, Rajimehr R, Kim BW, Ekstrom LB, Vanduffel W, Tootell RBH. 2006. The radial bias: a different slant on visual orientation sensitivity in human and nonhuman primates. Neuron. 51:661–670.

Schmahmann JD, Pandya D. 2006. Fiber pathways of the brain. New York: Oxford Univ Press.

Schmahmann JD, Pandya DN, Wang R, Dai G, D’Arceuil HE, de Crespigny AJ, Wedeen VJ. 2007. Association fibre pathways of the brain: parallel observations from diffusion spectrum imaging and autoradiography. Brain. 130:630–653.

Seehaus A, Roebroeck A, Bastiani M, Fonseca L, Bratzke H, Lori N, Vilanova A, Goebel R, Galuske R. 2015. Histological validation of high-resolution DTI in human post mortem tissue. Front Neuroanat. 9:98.

Sherbondy AJ, Dougherty RF, Ben-Shachar M, Napel S, Wandell BA. 2008a. ConTrack: finding the most likely pathways between brain regions using diffusion tractography. J Vis. 8:15 1–16.

Sherbondy AJ, Dougherty RF, Napel S, Wandell BA. 2008b. Identifying the human optic radiation using diffusion imaging and fiber tractography. J Vis. 8:12 1–12.

Silson EH, McKeefry DJ, Rodgers J, Gouws AD, Hymers M, Morland AB. 2013. Specialized and independent processing of orientation and shape in visual field maps LO1 and LO2. Nat Neurosci. 16:267–269.

Smith AT, Greenlee MW, Singh KD, Kraemer FM, Hennig J. 1998. The processing of first- and second-order motion in human visual cortex assessed by functional magnetic resonance imaging (fMRI). J Neurosci. 18:3816–3830.

Sotiropoulos SN, Hernández-Fernández M, Vu AT, Andersson JL, Moeller S, Yacoub E, Lenglet C, Ugurbil K, Behrens TEJ, Jbabdi S. 2016. Fusion in diffusion MRI for improved fibre orientation estimation: An application to the 3T and 7T data of the Human Connectome Project. Neuroimage. 134:396–409.

Sotiropoulos SN, Jbabdi S, Xu J, Andersson JL, Moeller S, Auerbach EJ, Glasser MF, Hernandez M, Sapiro G, Jenkinson M, Feinberg DA, Yacoub E, Lenglet C, Van Essen DC, Ugurbil K, Behrens TE, Consortium, W. U-Minn HCP. 2013. Advances in diffusion MRI acquisition and processing in the Human Connectome Project. Neuroimage. 80:125–143.

Takemura H, Ashida H, Amano K, Kitaoka A, Murakami I. 2012. Neural correlates of induced motion perception in the human brain. J Neurosci. 32:14344–14354.

Takemura H, Caiafa CF, Wandell BA, Pestilli F. 2016a. Ensemble Tractography. PLoS Comput Biol. 12:e1004692.

Takemura H, Rokem A, Winawer J, Yeatman JD, Wandell BA, Pestilli F. 2016b. A major human white-matter pathway between dorsal and ventral visual cortex. Cereb Cortex. 26:2205–14.

Thiebaut de Schotten M, Dell’Acqua F, Forkel SJ, Simmons A, Vergani F, Murphy DG, Catani M. 2011. A lateralized brain network for visuospatial attention. Nat Neurosci. 14:1245–1246.

Thomas C, Ye FQ, Irfanoglu MO, Modi P, Saleem KS, Leopold DA, Pierpaoli C. 2014. Anatomical accuracy of brain connections derived from diffusion MRI tractography is inherently limited. Proc Natl Acad Sci U S A. 111:46.

Tootell RB, Mendola JD, Hadjikhani NK, Ledden PJ, Liu AK, Reppas JB, Sereno MI, Dale AM. 1997. Functional analysis of V3A and related areas in human visual cortex. J Neurosci. 17:7060–7078.

Tootell RB, Tsao D, Vanduffel W. 2003. Neuroimaging weighs in: humans meet macaques in “primate” visual cortex. J Neurosci. 23:3981–3989.

Tournier JD, Calamante F, Connelly A. 2012. MRtrix: Diffusion tractography in crossing fiber regions. Int J Imaging Syst Technol. 22:53–66.

Tsao DY, Moeller S, Freiwald WA. 2008. Comparing face patch systems in macaques and humans. Proc Natl Acad Sci U S A. 105:19514–19519.

Tsao DY, Vanduffel W, Sasaki Y, Fize D, Knutsen TA, Mandeville JB, Wald LL, Dale AM, Rosen BR, Van Essen DC, Livingstone MS, Orban GA, Tootell RB. 2003. Stereopsis activates V3A and caudal intraparietal areas in macaques and humans. Neuron. 39:555–568.

Ungerleider LG, Desimone R. 1986. Cortical connections of visual area MT in the macaque. J Comp Neurol. 248:190–222.

Ungerleider LG, Galkin TW, Desimone R, Gattass R. 2008. Cortical connections of area V4 in the macaque. Cereb Cortex. 18:477–499.

Ungerleider LG, Haxby JV. 1994. “What” and “where” in the human brain. Curr Opin Neurobiol. 4:157–165.

Ungerleider LG, Mishkin M. 1982. Two cortical visual systems. In: Ingle DJ, Goodale MA, Mansfield RJW, editors. The analysis of visual behavior. Cambridge, MA: MIT Press. p. 549–586.

van den Heuvel MP, Bullmore ET, Sporns O. 2016. Comparative connectomics. Trends Cogn Sci. 20:345–361.

Vanduffel W, Fize D, Mandeville JB, Nelissen K, Van Hecke P, Rosen BR, Tootell RB, Orban GA. 2001. Visual motion processing investigated using contrast agent-enhanced fMRI in awake behaving monkeys. Neuron. 32:565–577.

Vanduffel W, Zhu Q, Orban GA. 2014. Monkey cortex through fMRI glasses. Neuron. 83:533–550.

Van Essen DC, Smith SM, Barch DM, Behrens TE, Yacoub E, Ugurbil K, Consortium, W. U-Minn HCP. 2013. The WU-Minn Human Connectome Project: an overview. Neuroimage. 80:62–79.

Vu AT, Auerbach E, Lenglet C, Moeller S, Sotiropoulos SN, Jbabdi S, Andersson J, Yacoub E, Ugurbil K. 2015. High resolution whole brain diffusion imaging at 7T for the Human Connectome Project. Neuroimage. 122:318–331.

Wade A, Augath M, Logothetis N, Wandell B. 2008. fMRI measurements of color in macaque and human. J Vis. 8:6 1–19.

Wakana S, Jiang H, Nagae-Poetscher LM, van Zijl PC, Mori S. 2004. Fiber tract-based atlas of human white matter anatomy. Radiology. 230:77–87.

Wandell BA. 2016. Clarifying human white matter. Annu Rev Neurosci. [Epub ahead of print].

Wandell BA, Dumoulin SO, Brewer AA. 2007. Visual field maps in human cortex. Neuron. 56:366–383.

Wandell BA, Rokem A, Perry LM, Schaefer G, Dougherty RF. 2015. Data management to support reproducible research. arXiv [q-bioQM].

Wandell BA, Winawer J. 2011. Imaging retinotopic maps in the human brain. Vision Res. 51:718–737.

Wandell BA, Yeatman JD. 2013. Biological development of reading circuits. Curr Opin Neurobiol. 23:261–268.

Wang L, Mruczek REB, Arcaro MJ, Kastner S. 2015. Probabilistic maps of visual topography in human cortex. Cereb Cortex. 25:3911–3931.

Warner CE, Kwan WC, Wright D, Johnston LA, Egan GF, Bourne JA. 2015. Preservation of vision by the pulvinar following early-life primary visual cortex lesions. Curr Biol. 25:424–434.

Webster MJ, Bachevalier J, Ungerleider LG. 1994. Connections of inferior temporal areas TEO and TE with parietal and frontal cortex in macaque monkeys. Cereb Cortex. 4:470–483.

Weiner KS, Grill-Spector K. 2015. The evolution of face processing networks. Trends Cogn Sci. 19:240–241.

Weiner KS, Jonas J, Gomez J, Maillard L, Brissart H, Hossu G, Jacques C, Loftus D, Colnat-Coulbois S, Stigliani A, Barnett MA, Grill-Spector K, Rossion B. 2016a. The Face-processing network is resilient to focal resection of human visual cortex. J Neurosci. 36:8425–8440.

Weiner KS, Yeatman JD, Wandell BA. 2016b. The posterior arcuate fasciculus and the vertical occipital fasciculus. Cortex. [Epub ahead of print]

Wernicke C. 1881. Lehrbuch der Gehirnkrankheiten für Aerzte und Studirende. Kassel Theodor Fischer.

Winawer J, Horiguchi H, Sayres RA, Amano K, Wandell BA. 2010. Mapping hV4 and ventral occipital cortex: the venous eclipse. J Vis. 10(5): 1–12.

Winawer J, Witthoft N. 2015. Human V4 and ventral occipital retinotopic maps. Vis Neurosci. 32:E020.

Witthoft N, Nguyen ML, Golarai G, Larocque KF, Liberman A, Smith ME, Grill-Spector K. 2014. Where is human V4? Predicting the location of hV4 and VO1 from cortical folding. Cereb Cortex. 24: 2401–2408.

Yeatman JD, Dougherty RF, Ben-Shachar M, Wandell BA. 2012a. Development of white matter and reading skills. Proc Natl Acad Sci U S A. 109:E3045–E3053.

Yeatman JD, Dougherty RF, Myall NJ, Wandell BA, Feldman HM. 2012b. Tract profiles of white matter properties: automating fiber-tract quantification. PLoS One. 7:e49790.

Yeatman JD, Rauschecker AM, Wandell BA. 2013. Anatomy of the visual word form area: adjacent cortical circuits and long-range white matter connections. Brain Lang. 125:146–155.

Yeatman JD, Weiner KS, Pestilli F, Rokem A, Mezer A, Wandell BA. 2014. The vertical occipital fasciculus: A century of controversy resolved by in vivo measurements. Proc Natl Acad Sci U S A. 111:E5214–E5223.

Yendiki A, Panneck P, Srinivasan P, Stevens A, Zöllei L, Augustinack J, Wang R, Salat D, Ehrlich S, Behrens T, Jbabdi S, Gollub R, Fischl B. 2011. Automated probabilistic reconstruction of white-matter pathways in health and disease using an atlas of the underlying anatomy. Front Neuroinform. 5:23.

Zarco W, Merchant H, Prado L, Mendez JC. 2009. Subsecond timing in primates: comparison of interval production between human subjects and rhesus monkeys. J Neurophysiol. 102:3191–3202.

